# Integrating genome-wide neutral and putatively adaptive variation\ resolves population structure and differentiation in a recently diverged albatross complex

**DOI:** 10.64898/2026.09.04.749127

**Authors:** Imogen Foote, Tom Oosting, Kath Walker, Graeme Elliott, Kalinka Rexer-Huber, Graham C Parker, Geoffrey K Chambers, Peter A Ritchie

## Abstract

Understanding patterns of evolution and divergence in populations is important for defining conservation units and informing taxonomy. However, when populations have low genetic diversity, or are closely related, genetic differentiation can be difficult to detect, particularly when relying on small numbers of genetic markers. Whole-genome sequencing allows for genome-wide identification of both neutral and outlier variation, improving the resolution of subtle population structure and providing insight into evolutionary processes. Here, we investigated genomic differentiation in a recently diverged and highly threatened albatross complex, the Antipodean (*D. antipodensis antipodensis*) and Gibson’s albatross (*D. a. gibsoni*), using genome-wide neutral and outlier datasets. Whole-genome resequencing of 86 individuals sampled across Antipodes Island (*D. a. antipodensis*) and the Auckland Islands (*D. a. gibsoni*) identified 381,176 neutrally evolving and 57 independently segregating outlier (putatively adaptive) SNPs. Analyses of both datasets revealed significant genetic differentiation between the Antipodean and Gibson’s albatross, no evidence of contemporary gene flow and evidence of selective sweeps suggesting local adaptation. Within-population structure was also identified for the Gibson’s albatross, with genetic differentiation among sample sites from different islands (Adams Island and Disappointment Island). Patterns of heterogenous differentiation across the genome suggest the taxa are on different evolutionary trajectories. Together with existing morphological and behavioural evidence, these genomic results support reassessment of their conservation and taxonomic status. More broadly, this study demonstrates the value of combining neutral and putatively adaptive genomic variation to resolve subtle population structure in recently diverged taxa.

## Introduction

Understanding how populations diverge into independently evolving lineages is a central goal of evolutionary biology. Accurate delineation of taxa is crucial for informing taxonomy and conservation, particularly in the face of current rapid and unprecedented species declines (Stanton et al., 2019; Theissinger et al., 2023). The advent of molecular data have transformed our ability to infer evolutionary relationships and demographic histories, and to quantify genetic variation within and between populations across the tree of life (Aspi et al., 2006; Baker et al., 1995; Hebert et al., 2004; Hedges & Poling, 1999; Plazzi et al., 2011). However, detecting genetic differentiation among recently diverged taxa remains challenging because relatively little genetic divergence may have accumulated and traditional marker datasets often lack the power to resolve subtle population structure (McCartney-Melstad et al., 2018; Spinks et al., 2014; Vendrami et al., 2017). Over the past decade, advancements in whole-genome sequencing have enabled population structure to be investigated with increasingly fine-scale resolution (Fang et al., 2022; Jospin et al., 2023; Martin et al., 2018), improving our ability to characterise evolutionary divergence and delineate recently diverged taxa.

As well as increased resolution, an advantage of whole-genome data is that genetic variation can be assessed across the entire genome, rather than just a small number of loci. Population divergence is shaped by genetic drift, selection and gene flow acting on genetic variation generated by mutation and recombination. These evolutionary mechanisms can act differently across the genome, leaving different genomic signatures (Beaumont & Balding, 2004; Storz, 2005; Vitti et al., 2013). Assessment of evolutionarily neutral loci can provide insight into demographic processes, including genetic drift and gene flow (Bertola et al., 2023; Fang et al., 2022; McCartney-Melstad et al., 2018). However, in the early stages of speciation, genomic differentiation at neutral markers may remain subtle as differences accumulate gradually. Whole-genome data also allow the identification of regions with elevated divergence (i.e., outlier loci). These outlier regions are candidates for divergent selection providing insights into the potential role of local adaptation during divergence (Allendorf et al., 2010; Kirk & Freeland, 2011). In some cases, assessment of outlier loci has revealed high genomic divergence where neutral loci showed genetic homogeneity or very weak differentiation (Berg et al., 2016; Milano et al., 2014; Tigano et al., 2017). Thus, integrating assessment of genome-wide neutral variation with outlier loci is critical to gain a more complete understanding of evolutionary history and divergence, and improves the evidence available for delineating conservation units and informing taxonomy (Funk et al., 2012; Kirk & Freeland, 2011; Stanton et al., 2019).

Albatrosses (family Diomedeidae) provide an excellent model to explore the power of genomic data for resolving evolutionary patterns and subtle population structure in recently diverged lineages. The Antipodean and Gibson’s albatrosses (*Diomedea antipodensis antipodensis* and *D. a. gibsoni*, respectively) are members of the wandering albatross complex endemic to Aotearoa/New Zealand (NZ), with breeding restricted to small offshore islands in NZ’s subantarctic region (Figure 1). Both populations have suffered steep declines in recent decades, largely due to mortality from interaction with longline fishing vessels (Elliott et al., 2025; Rexer-Huber et al., 2025; Richard et al., 2024). Therefore, accurate definition of conservation units is required, yet questions regarding their taxonomy and degree of population divergence remain (Foote et al., 2025). The taxonomic status of the Antipodean and Gibson’s albatrosses has been the subject of numerous revisions (Burg & Croxall, 2004; Robertson & Nunn, 1998; Robertson & Warham, 1992). Previous genetic work using microsatellite and mitochondrial markers revealed among-population structure, but overall low genetic differentiation between the populations at these markers, prompting reclassification from species to subspecies (ACAP, 2006; Burg & Croxall, 2004). In contrast, non-genetic data suggest some degree of divergence between the taxa, including morphological and plumage differences (Walker et al., unpublished data, 2024), asynchronous breeding (Walker & Elliott, 2005) and distinct foraging distributions (Walker & Elliott, 2006). Variation in adult survival, productivity and recruitment between the subspecies (Parker et al., 2023; Rexer-Huber et al., 2020; Walker et al., 2023) also indicates a lack of demographic connectivity (Ovenden, 2013). Together, these data highlight the need for a genome-wide reassessment of evolutionary divergence. Such an assessment provides an opportunity to evaluate the utility of combining neutral and putatively adaptive variation for improving inference of population differentiation, conservation unit delineation and taxonomy.

**Figure 1.**
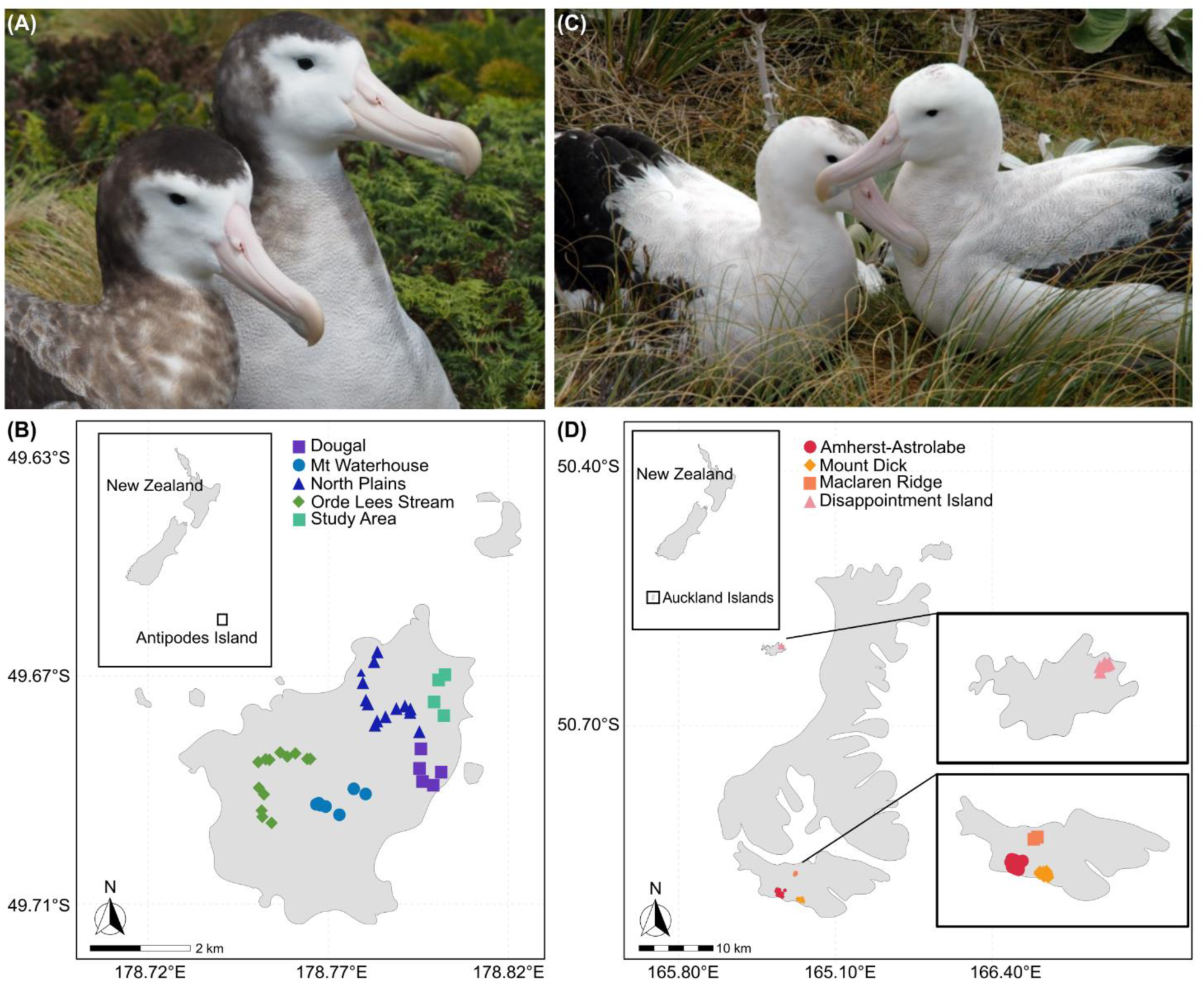
**A)** A female (left) and male (right) Antipodean albatross (*Diomedea antipodensis antipodensis*) breeding pair, **B)** map of Antipodes Island showing the main breeding location and sampling locations of 43 breeding adult Antipodean albatrosses, **C)** a female (left) and male (right) Gibson’s albatross (*D. a. gibsoni*) breeding pair and **D)** map of the Auckland Islands showing the main breeding location (Adams Island – bottom) and the sampling locations of 43 breeding adult Gibson’s albatross across Adams Island and Disappointment Island. Inset boxes show location of islands relative to NZ mainland (top left) and magnification of sampling sites (bottom right). The distance between Antipodes Island and the Auckland Islands is approximately 900 km. *Photos: Kath Walker.* **Note:** “Study Area” is the name of a sampling location where a long-term population monitoring programme has been undertaken since 1994.

The aim of this study was to reassess population structure and genomic differentiation between Antipodean and Gibson’s albatrosses using a whole-genome dataset. Specifically, we quantified genome-wide differentiation at neutral and outlier (putatively adaptive) loci. By integrating genome-wide neutral and outlier variation, we present a more informative framework for assessing evolutionary divergence and informing conservation unit delineation and taxonomy in recently diverged taxa.

## Materials & Methods

### Sampling, DNA extraction and sequencing

Blood samples were collected from 86 nesting adult individuals from two breeding colonies during the 2021/2022 breeding season; 43 Antipodean (*D. a. antipodensis*) samples from Antipodes Island and 43 Gibson’s (*D. a. gibsoni*) albatross samples from the Auckland Islands (35 from Adams Island and 8 from Disappointment Island; Figure 1). Sampling methods were approved by the Victoria University of Wellington Animal Ethics Committee (VUW AEC 30055). Samples were collected across a broad geographic range for each subspecies with an even sampling ratio of males and females. As males are the non-dispersing sex, a sampling protocol was followed in which only males nesting more than 200 m apart were sampled to minimise the chance of sampling closely related individuals. Blood was preserved in Queen’s lysis buffer (10 mM Tris, 10 mM NaCl, 10 mM EDTA, 1% n- lauroylsarcosine, pH 7.5; Seutin et al., 1991), and stored at 4°C.

DNA was extracted using the QIAGEN DNeasy Blood & Tissue Kit (QIAGEN, Inc.). Purified DNA was stored in 10 mM Tris-HCl pH 8 and 0.1 mM EDTA and stored at 4°C. Library preparation and sequencing were performed by the Australian Genome Research Facility (AGRF). Libraries were prepared with the Illumina DNA PCR-Free Prep and sequencing was performed using an Illumina NovaSeq 6000 S4, to produce 150 bp paired-end reads.

### Read alignment and genotyping

Read quality was assessed using FastQC v0.11.7 (Andrews, 2010), and MultiQC v1.7 (Ewels et al., 2016). High-quality genome assemblies are available for both taxa (Foote et al., 2026) and analyses were undertaken with reads mapped to each reference. However, both reference genomes produced very similar results, so only analyses using the Antipodean albatross reference genome are presented (see Supplementary Materials B for Gibson’s albatross reference results). Mapping was performed using the Paleomix v1.3.7 pipeline which performs preprocessing steps and aligns reads to the reference genome (Schubert et al., 2014). First, adapter trimming is performed with AdapterRemoval v2.3.3 (Schubert et al., 2016) to remove Illumina sequencing adapters, allowing a 33% mismatch rate (--mm: 3) and minimum read length of 25 bp (--minlength: 25). Reads were mapped with the Burrows-Wheeler Aligner (BWA) v0.7.17 bwa-mem algorithm, filtering reads with a mapping quality below 25 (--MinQuality 25; Li & Durbin, 2009). Reads that only partially aligned to the reference genome (soft-clipped reads) were filtered out of the bam file and a new index file was created using SAMtools view and index (Danecek et al., 2021). Following read mapping, two individuals with much higher average sequencing depth were randomly subsampled down to the mean depth using SAMtools view.

Variant calling was performed with BCFtools v1.19 mpileup and call commands to produce VCF output for both subspecies (Danecek et al., 2021). Genotyping was performed on the largest 65 scaffolds only, as over 95% of each genome was contained within these scaffolds (Foote et al., 2026). Missing annotations were added to the VCF using the BCFtools +fill-tags plugin. The final VCF (AllSites dataset) contained both variant and invariant sites to enable unbiased calculation of genome diversity statistics such as π and Watterson’s θ (Korunes & Samuk, 2021). The AllSites dataset was filtered to create an additional dataset retaining only variant sites (VariableSites dataset) for further population genomic analysis.

### SNP quality filtering

Depth, missingness, site quality and allele frequency statistics of both datasets were calculated using VCFtools v0.1.15 (Danecek et al., 2011) to assess the characteristics of the data and determine appropriate thresholds for filtering (Figure SA1). The impact of different minor allele frequency (MAF) and missingness filtering thresholds on data analysis was tested, however the observed population structure did not appear sensitive to variation in these thresholds (Figure SA2). For the AllSites dataset, filtering was performed using the workflow recommended by Korunes and Samuk (2021). Briefly, filtering was performed only on variant sites, as lower overall quality of invariant sites can lead to significant data loss. Filtering was performed using VCFtools to remove low-quality SNPs (--minQ 30, -- max-missing 0.95, --min-meanDP 10, --max-meanDP 30) and to remove indels (--remove-indels). For the VariableSites dataset, filtering was performed as follows: VCFtools v0.1.15 was used for filtering to set all individual genotypes with low allelic depth to missing (-- minDP 3) and remove indels (--remove-indels). Further filtering was performed to remove low-quality SNPs (--minQ 30, --max-missing 0.95, --maf 0.05, --min-meanDP 8, --max-meanDP 30) and retain only biallelic SNPs (--min-alleles 2, --max-alleles 2). A custom R script adapted from Pinsky et al. (2021) was used to identify SNPs suffering from allelic imbalance, however no sites were identified for removal. This script performs a two-sided binomial test to evaluate whether there is a significant deviation from an expected equal distribution of reference and alternative alleles within individuals at heterozygous sites. The filtered AllSites and VariableSites SNP datasets were converted from VCF to both GDS and PLINK file format for further steps using the *SNPRelate* package (Zheng et al., 2012) and a custom R script.

### SNP neutral and outlier filtering

Identification of outlier loci was performed using pcadapt (Luu et al., 2017) and BayPass (which accounts for existing population structure; Gautier, 2015) on the VariableSites data. First, pcadapt was run with K=10 and the resulting screeplot was used to determine the number of informative principal components using Cattell’s rule. The programme was rerun with K=1, with a minimum allele frequency of 0.05 (min.maf 0.05). Baypass was run on both the real data and a simulated dataset generated using the Baypass R function *simulate.baypass()*. An appropriate XtX statistic threshold for identifying outlier loci was determined using the simulated data and then applied to filtering the output from the real data. Overlapping outlier SNPs identified by both BayPass and pcadapt were retained for subsequent outlier analysis (Outlier dataset) and SNPs in linkage disequilibrium were removed using PLINK v1.09 (--indep-pairwise 50 5 0.2; Purcell et al., 2007).

All outlier loci identified in the previous step were removed from the full VariableSites dataset using VCFtools (--exclude-positions) to retain a neutral SNP dataset (Neutral dataset). SNPs deviating from Hardy-Weinberg equilibrium (-hwe 0.001) were identified per population (following the “Out All” approach suggested by Pearman et al. (2022)) and removed using VCFtools. Finally, SNPs in linkage disequilibrium were removed using PLINK v1.09 (--indep-pairwise 50 5 0.2; Purcell et al., 2007). The *vcfrandomsample* function of vcflib v1.0.1 was used to create a random subset of ∼10,000 SNPs (Subset dataset) for programmes which do not perform well with large datasets (e.g., BA3-SNPs, NeEstimator2).

### Genetic diversity and population structure

Nucleotide diversity statistics including π, D_XY_ and Watterson’s θ were estimated from the AllSites dataset using the pixy software v2.0.0 (Korunes & Samuk, 2021) with 10,000 bp window sizes (--window_size 10000). Genome-wide averages were calculated as per the author’s guidelines. All subsequent analyses were performed on both the Neutral and Outliers datasets. Observed (*H*_o_) and expected heterozygosity (*H*_e_) were calculated with the function *gl.report.heterozygosity()* in the *dartR* package (Gruber et al., 2018). Pairwise weighted *F_ST_* values were calculated between subspecies and sampling sites for each SNP to assess population differentiation using the Weir & Cockerham, 1984 method implemented with the snpgdsFST function in SNPRelate (Weir & Cockerham, 1984; Zheng et al., 2012). Bootstrapping with 10,000 iterations was performed to calculate 95% confidence intervals. A hierarchical Analysis of Molecular Variance (AMOVA) was performed using the R package *poppr* (Kamvar et al., 2014) to compare the genetic variation between subspecies. Significance-levels were determined using the randtest function from the *ade4* package in R (Chessel et al., 2009). PCAs were performed based on a Genetic Relationship Matrix (GRM) using *SNPRelate* in R (Zheng et al., 2012). ADMIXTURE was run using the Neutral dataset only for values of K from 1 to 5 to determine the number of K ancestral populations with the lowest cross-validation error, which was then rerun with 10,000 iterations (Alexander et al., 2009).

### Estimation of effective population size, population demographic history and gene flow

Effective population sizes (N_e_) were estimated from the Subset dataset for each subspecies with NeEstimator2 (Do et al., 2014) using the Linkage Disequilibrium method and monogamy mating model (Waples & Do, 2008). Upper and lower confidence intervals for N_e_ estimates were calculated based on jackknife resampling. Recent demographic history was inferred using the LD-based method in GONE with default options except analysis was restricted to the first 40 (largest) scaffolds because scaffolds with a small number of SNPs can cause issues with analysis in GONE (Santiago et al., 2020). A constant recombination rate of 1 cM per Mb was assumed according to the developer’s recommendation when no genetic map is available. Analysis was performed separately for the Antipodean and Gibson’s subspecies as the programme assumes a closed population without recent admixture.

Contemporary gene flow between the populations was assessed with BA3-SNPs (Mussmann et al., 2019), a modification of BayesAss (Wilson & Rannala, 2003) to utilise large SNP datasets. First, the BA3-SNPs-autotune script was used with the Subset dataset to automatically determine the mixing parameters for migration rates (-m), allele frequencies (- a) and inbreeding coefficients (-f). These values were used to run BA3-SNPs with 15 million iterations (-i15000000) and a burn-in period of 1,500,000 iterations (-b1500000). The programme was run twice; first between subspecies (Antipodean and Gibson’s albatross), and then between sampling island locations (Antipodes Island, Adams Island and Disappointment Island).

### Detection of selective sweeps

To further investigate signatures of selection in the genome, Cross-Population number Segregating sites by Length (XP-nSL) statistics were calculated using Selscan v3.0.1 (Rahman et al., 2026). This statistic compares haplotype patterns between two populations to detect regions of Extended Haplotype Homozygosity (EHH) in one population indicative of recent local selective sweeps. XP-nSL is preferred over the related XP-EHH statistic for unphased datasets and when no genomic map is available (Ferrer-Admetlla et al., 2014; Szpiech et al., 2021). XP-nSL was calculated with Antipodean individuals as population 1 (--vcf) and Gibson’s individuals as population 2 (--vcf-ref) and with options --max-gap 200000, --max-extend-nsl 500000, --unphased and --xpnsl. XP-nSL scores were normalised against the genome-wide distribution using the selscan norm command with the options --xpnsl, -- qbins 10, --win-size 10000 and --bp-win.

The most extreme value per 10 kb window was selected for plotting, and the most extreme 1% of windows containing at least 10 SNPs were identified, as well as windows containing outlier SNPs identified by Baypass and pcadapt. These windows were annotated to investigate their potential function by extracting overlapping genes and associated Gene Ontology (GO) terms (where available) from the available genome annotation (Foote et al., 2026).

## Results

### Data quality and filtering

In total, 13.8 billion reads were produced, giving an average of 160 million reads or 20x sequencing depth per individual. Reads from all individuals of both subspecies were of high quality (Phred score > 30; Figure SA3).

During genotyping, 1,225,175,514 sites (All Sites) were identified across the genome including 7,285,599 polymorphic sites (Variable Sites), with an average depth of 13.3x. After quality filtering, 1,150,970,116 sites were retained in the AllSites dataset, and 2,400,435 polymorphic SNPs retained in the VariableSites dataset. BayPass identified 19,068 outlier SNPs, and pcadapt identified 6,672 outlier SNPs. Of these, there were 746 overlapping SNPs. Following LD pruning, 57 independently segregating outlier SNPs were retained for further population genomic analysis. Following removal of all identified outlier SNPs from the high-quality SNP dataset and pruning for linked SNPs, the final dataset contained 381,176 neutrally evolving and independently segregating SNPs. No individual samples were excluded from further analysis based on quality metrics (Figure SA4).

### Genetic diversity and population structure

Estimates of nucleotide diversity, as well as heterozygosity were consistent and similar across both taxa (Table 1). When assessing the neutral SNP dataset, AMOVA showed that most observed variation was contained within samples (∼99%) rather than between subspecies (∼4%) or between samples within subspecies (∼-3%; this total variation summed to greater than 100% due to a slightly negative covariance estimated between samples, which can occur in the absence of genetic structure within populations). However, only the between-subspecies differentiation was significant (p = 0.001, compared with p = 0.929 and p = 0.660, respectively; Table SA1). The outlier SNP dataset showed that much more genetic variation was observed between subspecies (59.2%, *p* = 0.001) than between samples within subspecies (8.3%, *p* = 0.001) and within samples (32.5%, p = 0.001), with all comparisons statistically significant (Table SA2).

**Table 1.** Genetic diversity statistics calculated using a dataset of 381,176 neutral SNPs or a genome-wide SNP dataset (including invariant sites) for the Antipodean and Gibson’s albatross. *H*_o_ = observed heterozysity (SE), *H*_e_ = expected heterozygosity (SE), *π* = average nucleotide diversity, *θ* = average Watterson’s estimator, *F_IS_* = inbreeding coefficient (1 − *H*_o_/*H*_e_), *D_XY_* = average absolute nucleotide divergence, *F_ST_*= average genetic differentiation. †Calculated from AllSites dataset, *p<0.001.

| | N | $H_o$ | $H_e$ | $\pi$† | $\theta$† | $F_{IS}$ | $D_{XY}$† | $F_{ST}$ |
| --- | --- | --- | --- | --- | --- | --- | --- | --- |
| Antipodean | 43 | 0.311<br>(0.0003) | 0.300<br>(0.0003) | 0.0009 | 0.0009 | -0.037 | 0.0009 | 0.037* |
| Gibson's | 43 | 0.312<br>(0.0003) | 0.298<br>(0.0003) | 0.0009 | 0.0008 | -0.047 |  |  |

Population structure analyses using a range of methods identified generally similar results from the Neutral dataset, revealing two distinct genetic groups corresponding to the two taxa (Antipodean and Gibson’s albatross). PCA results support the separation of two distinct clusters. Principal component 1 (PC1) accounted for 5.16% of the total genetic variation and clear separation of Antipodean individuals from Gibson’s individuals. PC2 explained 1.55% of the observed variation and showed separation of individuals from the Disappointment Island sampling site from the rest of the Gibson’s albatross samples (Figure 2A). *D_XY_* and *F_ST_* estimates revealed a low, but significant, level of differentiation between Antipodean and Gibson’s albatross (Table 1). Pairwise comparisons revealed very low to no differentiation between different sampling sites within taxa, with the largest level of within-taxon differentiation being observed between Disappointment Island and the rest of the Gibson’s albatross sites (*F_ST_* ≈ 0.01; Figure SA5). ADMIXTURE results showed the lowest cross- validation error was obtained with K = 2 (Table SA3), which corresponded to the Antipodean and Gibson’s albatross subspecies (Figure 2B).

**Figure 2.**
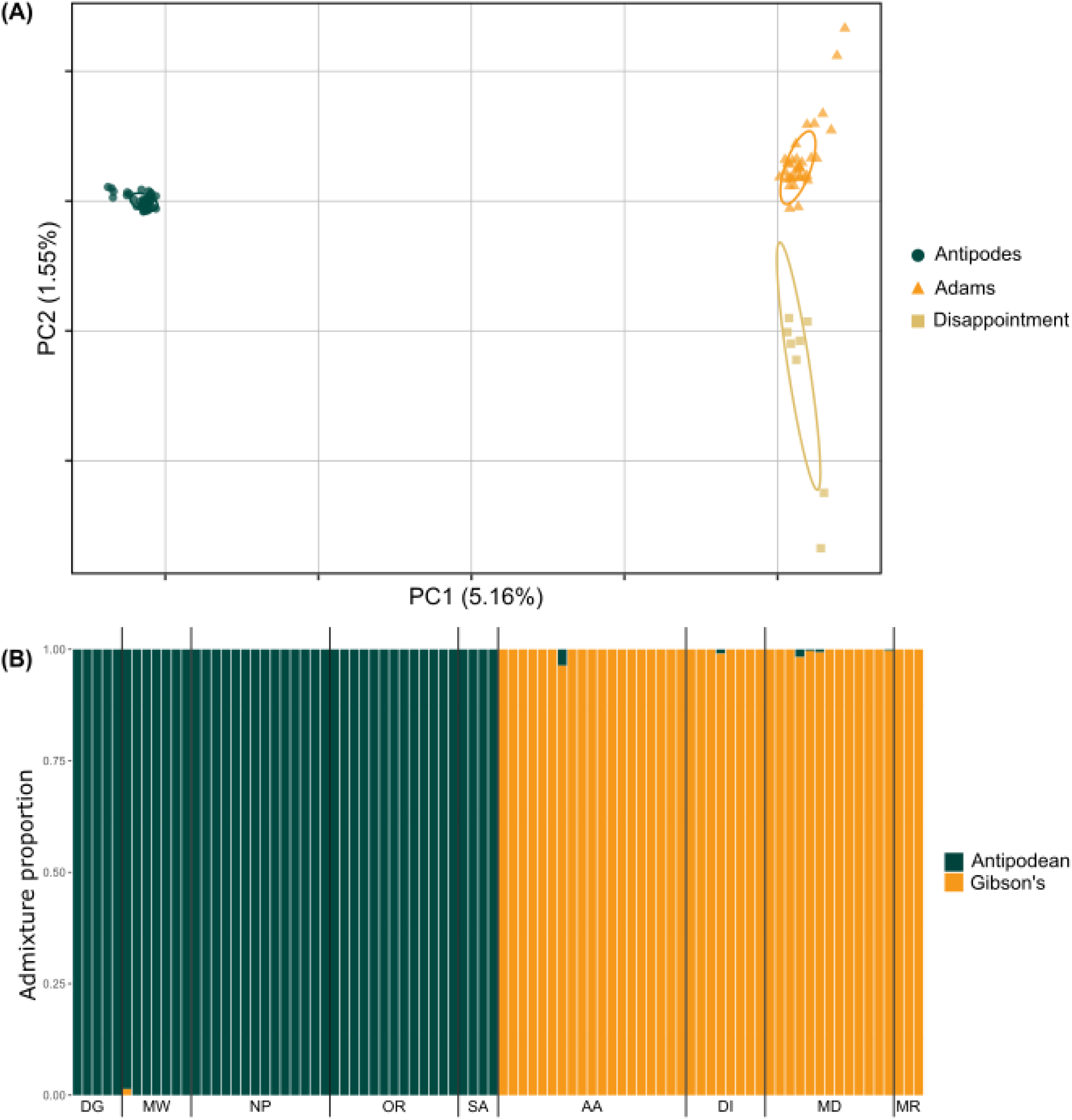
**A)** Principal Component Analysis (PCA) based on 381,176 neutral SNPs showing the genetic clustering of Antipodean albatross (left cluster) and Gibson’s albatross (right cluster). Samples are grouped by sampling island. **B)** ADMIXTURE plot for K = 2 showing the ancestry proportions of individuals from Antipodean and Gibson’s albatross populations using 381,176 neutral SNPs. Solid black lines indicate groupings of individuals from each sampling site. DG = Dougal, MW = Mt Waterhouse, NP = North Plains, OR = Orde Lees Stream, SA = Study Area, AA = Amherst Amsterdam, DI = Disappointment Island, MD = Mt Dick, MR = Maclaren Ridge

### Effective population size, demographic history and gene flow

Contemporary gene flow analysis with BA3-SNPs suggested no significant migration between the Antipodean and Gibson’s subspecies over the past few generations (not significantly different from zero), however recent migration was observed from Adams Island to Disappointment Island (*m* = 0.2727; Figure 3A, Table SA4). The Antipodean albatross had a larger contemporary effective population size (N_e_ = 2,258; 1,043 – ∞ 95% CI) than the Gibson’s albatross (N_e_ = 1,157; 598 – 11,669 95% CI; Table SA5). Recent demographic analysis with GONE also indicated a higher N_e_ of the Antipodean subspecies over the past 200 generations, with a rapid recent population expansion and decline, while the Gibson’s albatross has mostly remained small with gradual declines over time (Figure 3B).

**Figure 3.**
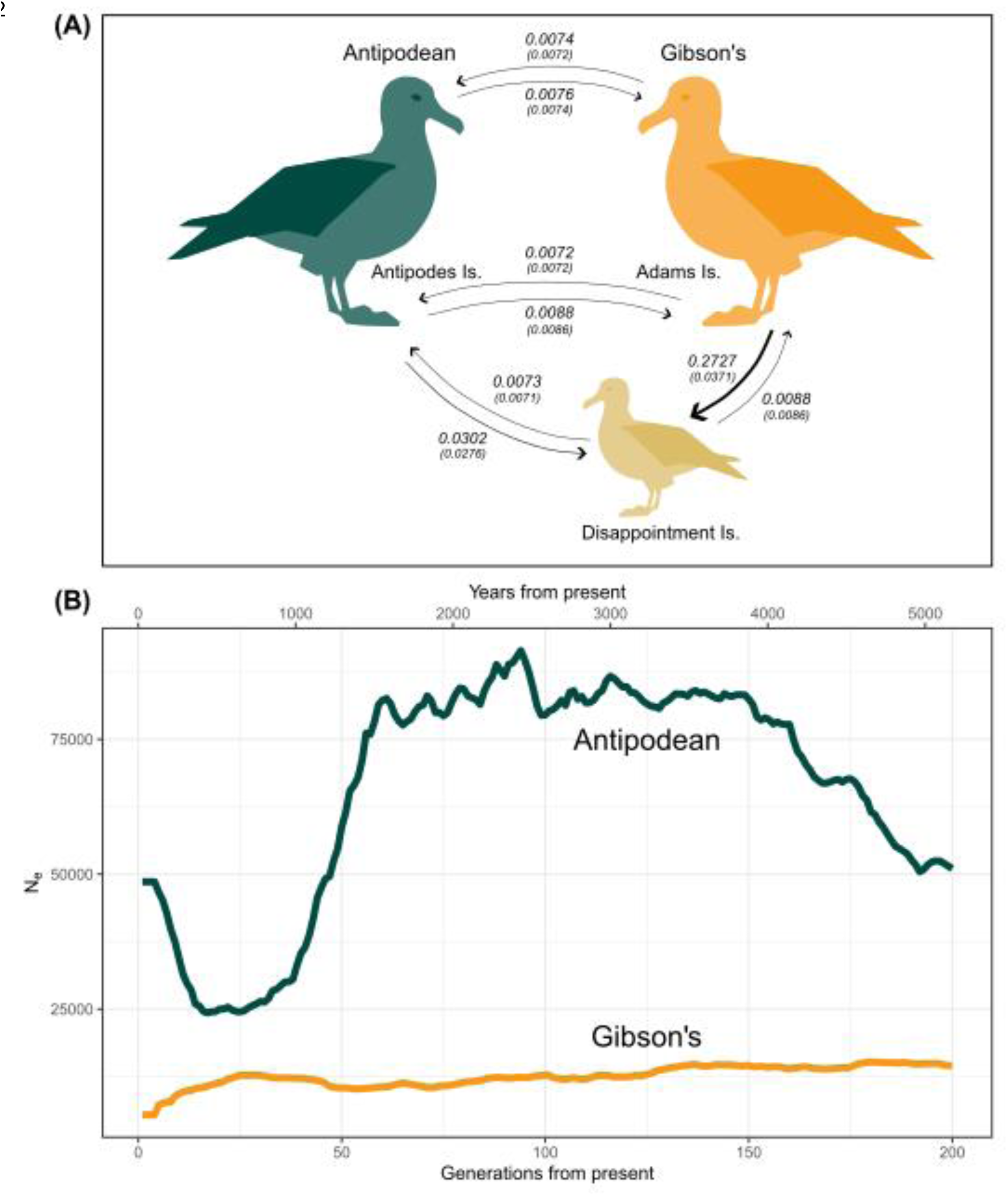
**A)** Recent migration rates between populations (Antipodean and Gibson’s albatross) and islands (Antipodean = Antipodes Island, Gibson’s = Adams Island and Disappointment Island) estimated with BA3-SNPs. Arrows show direction and estimate of the mean (SD) rate of inferred migration. Values indicate the proportion of individuals in the receiving population that are migrants from the source population. **B)** Recent demographic history of the Antipodean and Gibson’s albatross inferred using GONE. N_e_ estimates are averages of estimates over 40 independent iterations of the programme. Years from present is calculated using an estimated generation length of 25.86 years provided by Bird et al. (2020).

### Population differentiation – outlier loci

When assessing the dataset of 57 independent outlier loci, PCA results support the separation of two genetic clusters (Figure 4). Principal component 1 (PC1) accounted for 50.98% of the total genetic variation and clear separation of Antipodean individuals from Gibson’s individuals. PC2 explained 6.60% of the observed variation showing a wider spread of variation amongst the Gibson’s individuals with separate clustering of the Disappointment Island individuals, while the Antipodean individuals formed a relatively tight cluster. Estimates of heterozygosity for the Gibson’s albatross at outlier loci were slightly higher than at neutral loci and roughly double the estimated heterozygosity of Antipodean albatross at these loci (Table 2). Much higher genetic differentiation was observed between the taxa at outlier loci compared to neutral loci (F_ST_ = 0.509; Table 2).

**Figure 4.**
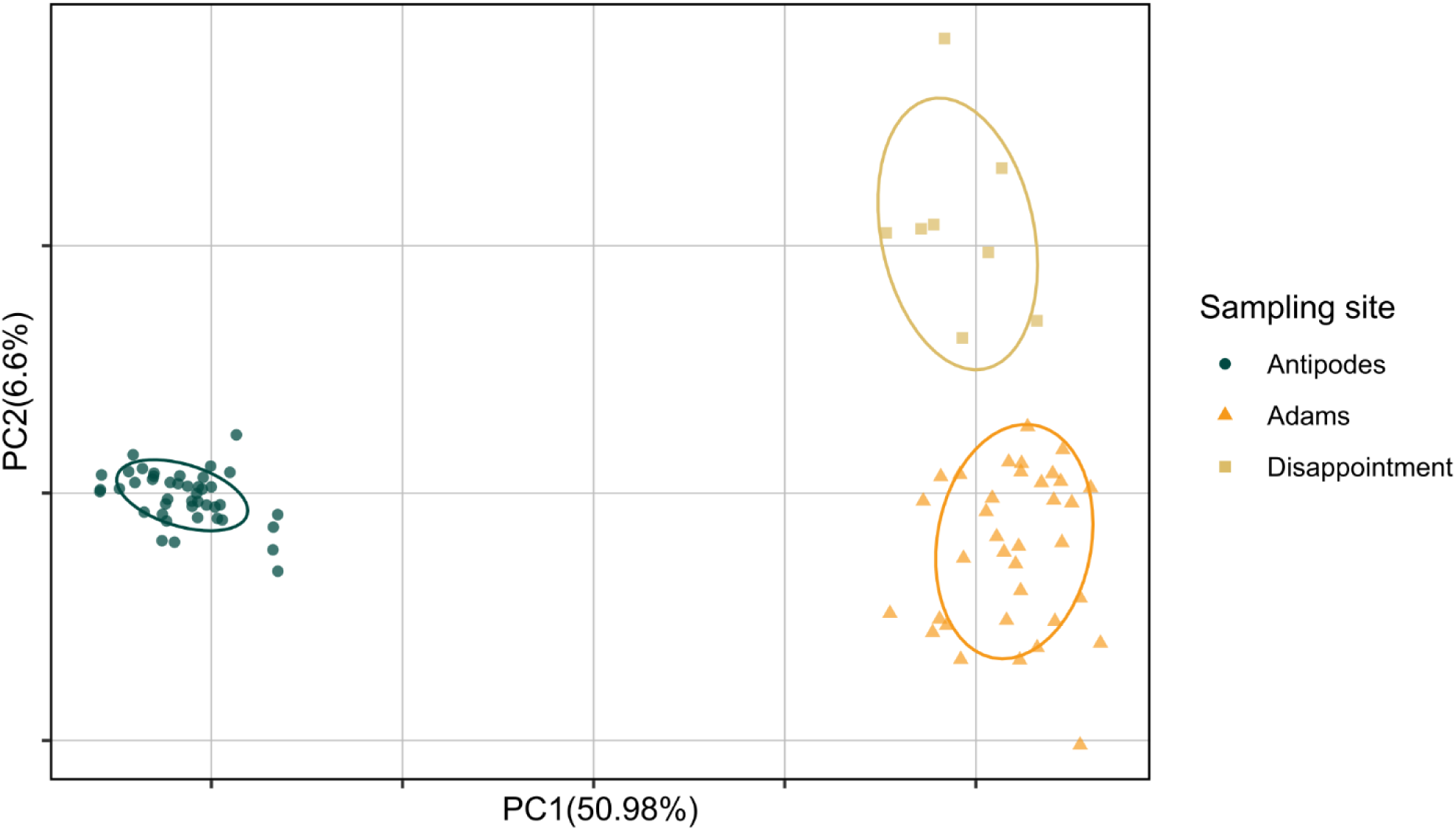
Principal Components Analysis (PCA) based on 57 outlier SNPs showing the genetic clustering of Antipodean albatross (left cluster) and Gibson’s albatross (right cluster). Samples are grouped by sampling island.

**Table 2.** Genetic diversity statistics calculated using a dataset of 57 outlier SNPs for the Antipodean and Gibson’s albatross. *H*_o_ = observed heterozysity (SE), *H*_e_ = expected heterozygosity (SE), *F_IS_* = inbreeding coefficient (1 − *H*_o_/*H*e), *F_ST_* = average genetic differentiation. *p<0.001.

| | N | $H_o$ | $H_e$ | $F_{IS}$ | $F_{ST}$ |
| --- | --- | --- | --- | --- | --- |
| Antipodean | 43 | 0.192<br>(0.0213) | 0.186<br>(0.0182) | -0.032 | 0.509* |
| Gibson's | 43 | 0.387<br>(0.0200) | 0.387<br>(0.0139) | 0.000 |  |

### Selection signatures

Loci displaying signatures of selection were identified across the genome with support from outlier analysis (pcadapt and Baypass) and haplotype homozygosity patterns (XP-nSL; Figure 5A). In total, 24 genes, were identified within windows showing strong XP-nSL signal and containing outlier SNPs (Supplementary Materials C). The top selection signatures were identified on scaffolds 6, 7 and 22, where elevated XP-nSL and a high density of outlier SNPs were observed. These regions display associated increased relative differentiation (F_ST_) between the Antipodean and Gibson’s albatross, however, did not consistently coincide with changes in absolute differentiation (D_XY_), Tajima’s D or π. (Figure 5B). An unusual pattern of high D_XY_, Tajima’s D and π was observed on scaffold 6 (a putative Z-chromosome scaffold; Foote et al., 2026) between ∼0 – 18 Mb. Further investigation suggested this unusual pattern was driven by high diversity and differentiation in females (Figure SA6). It is therefore hypothesised that this region represents an “evolutionary stratum”, resulting from recombination suppression in a formerly Pseudo Autosomal Region (PAR; Lahn & Page, 1999). The elevated π and D_XY_ in females is likely due to mis-mapping of a small proportion of divergent W-chromosome reads to this Z-chromosome scaffold. The genome assembly does not contain an assembled W-chromosome (Foote et al., 2026), potentially contributing to the mis-mapping of W-chromosome reads. This hypothesis is supported by increased sequencing depth observed for females (mean depth = 9.4x in this region compared to 7.7x for the rest of the scaffold; males mean depth = 14.7x across scaffold) and increased heterozygosity in females (otherwise expected to be zero) in this region (Figure SA7).

**Figure 5.**
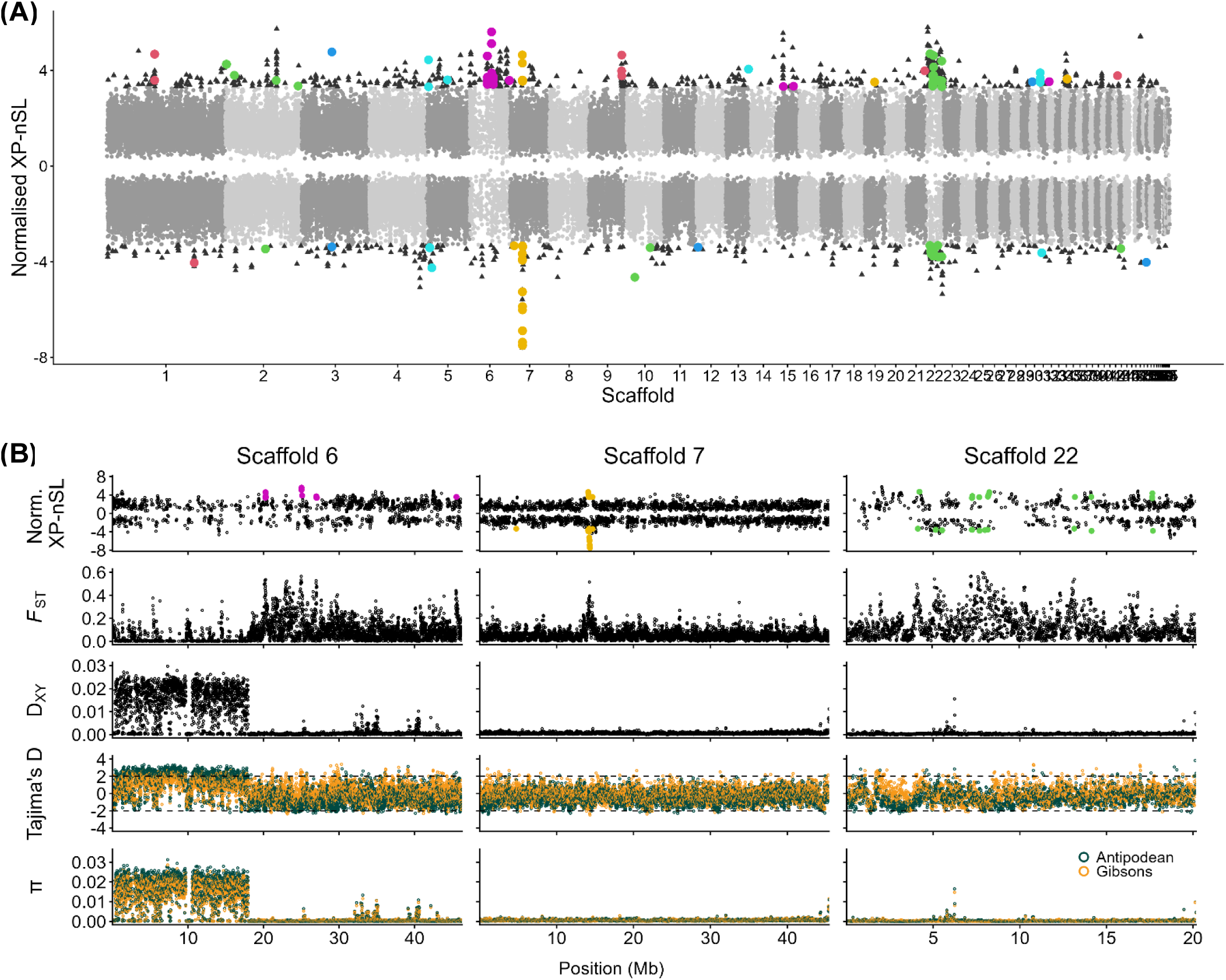
**A)** Manhattan plot of normalised XP-nSL values in 10 kilobase (kb) windows across the genome in Antipodean and Gibson’s albatrosses. Positive and negative XP-nSL values suggest evidence for a selective sweep in the Antipodean and Gibson’s populations, respectively. Plot shows only the most extreme value per window (light and dark grey points, coloured by scaffold), with the top 1% most extreme values represented by black triangles. Windows within the top 1% that contain outlier SNPs identified by Baypass and pcadapt are shown by coloured circles (coloured by scaffold). Names, genomic positions and putative functions of genes identified within these windows with evidence from XP-nSL, Baypass and pcadapt are provided in Supplementary Materials C. **B)** Normalised XP-nSL, F_ST_, D_XY_, Tajima’s D (dashed horizontal lines at +2 and −2 indicate approximate significance thresholds) and nucleotide diversity (π) across three scaffolds of interest. Coloured points in XP-nSL plots show windows with extreme values that also contain outlier SNPs identified by Baypass and pcadapt as per (A).

## Discussion

The results of this study revealed low but significant levels of genomic differentiation with no significant inferred gene flow between the Antipodean and Gibson’s albatross. The presence of these two distinct genetic clusters was apparent across all analyses of both neutral and outlier (putatively adaptive) SNP datasets and consistent regardless of the reference genome used for mapping reads (Antipodean vs. Gibson’s albatross reference individual). Combined evidence from outlier loci and XP-nSL statistics also identified regions of the genome showing signatures consistent with selection, suggesting the populations may be adapting to local conditions. This study presents the first population level whole-genome dataset for *Diomedea* albatrosses. Importantly, the findings demonstrate the power of integrating genome-wide neutral variation to reveal subtle population structure and resolve genetic connectivity, alongside assessment of putatively adaptative genomic regions to identify genomic regions where differentiation may be maintained by divergent selection, for distinguishing recently diverged taxa. These findings are discussed in relation to their implications for our understanding of albatross evolution and their significance for taxonomy and conservation.

### Genomic diversity and population structure

The average heterozygosity within each taxon was similar between the Antipodean and Gibson’s albatross (Antipodean H_e_ = 0.311; Gibson’s H_e_ = 0.312; Table 1) yet are lower than previous estimates calculated using a dataset of 9 microsatellite loci (Antipodean H_e_ = 0.43; Gibson’s H_e_ = 0.40; Burg & Croxall, 2004). This difference is not surprising as the previous genetic markers were likely selected for their levels of high variability (Brown et al., 1986; Väli et al., 2008). In comparison, the genomic data provide the most accurate representation of genetic diversity that current genetic methods allow. Although it is difficult to make inferences about comparisons of heterozygosity calculated using different datasets, similar levels of heterozygosity were observed in Northern Buller’s albatross (H_e_ = 0.32) and Southern Buller’s albatross (H_e_ = 0.29), estimated using genomic data (Wold et al., 2021).

Overall observed genetic diversity is low in both taxa, yet there is no evidence of any reduction in heterozygosity consistent with inbreeding (Table 1). This observation is consistent with findings of low genetic diversity in other *Diomedea* albatrosses, hypothesised to be inherited from a common ancestor nearly 1 Mya (Milot et al., 2007). Such low levels of diversity in these taxa may be the result of low mutation rates due to small long-term population sizes and low fecundity and does not appear to have had a detrimental impact on populations historically.

High concordance was observed across all analyses and datasets (neutral and outlier SNPs), showing two distinct groups corresponding to Antipodean and Gibson’s subspecies on Antipodes Island and the Auckland Islands, respectively. At neutral loci, genomic differentiation was low but significant (Table 1) while outlier data showed small regions of more substantial differentiation occurring across the genome (Table 2). Similar to the levels of heterozygosity, estimates of differentiation at neutral loci reported here between birds sampled on Antipodes Island and Adams Island (*F_ST_* = 0.040 without Disappointment Island samples; see Table **1** SA5), are lower than previously reported (mtDNA: *F_ST_* = 0.12, microsatellites: *F_ST_* = 0.07; Burg & Croxall, 2004) which is likely attributable to higher variability of selected markers in previous genetic studies in contrast to the current genome- wide view. Estimates of absolute divergence also show very low divergence between the taxa (D_XY_ = 0.0009) which may be explained by ongoing gene flow; however, we found no evidence for recent migration between the subspecies with BA3-SNPs and minimal admixture was observed with ADMIXTURE. This is consistent with lack of evidence of hybridisation between the taxa over three decades of monitoring and suggests that observed low levels of differentiation may be a result of retained ancestral variation due to recent divergence combined with low historical genetic diversity, rather than ongoing gene flow (Friesen et al., 2007). These findings, in combination with the observed structure, indicate that although most alleles may remain shared, drift and/or selection have generated consistent allele frequency differences between the taxa. Such differences have resulted in distinct genetic groupings, suggesting ecological or behavioural factors are acting as barriers to gene flow.

PCA analyses reveal that the genetic structure observed is best explained by geography, with each island displaying unique patterns of genetic diversity. As well as the clear separation of Antipodean and Gibson’s individuals, both neutral and outlier datasets revealed structuring within the Gibson’s albatross population, with slight segregation of the samples collected on Disappointment Island (DI) from the rest of the Gibson’s albatross samples collected on Adams Island (Figure 2A; Figure 1). Many seabird species in the Southern Ocean are wide-ranging and highly mobile, and there are no apparent barriers to gene flow between geographically distinct populations. Yet, population structuring is often observed (this study; Cristofari et al., 2019; Danckwerts et al., 2021; Rexer-Huber et al., 2019; Wold et al., 2021). Differences in at-sea distribution and strong philopatry are thought to drive population structure in many Southern Ocean seabirds (Friesen et al., 2007; Munro & Burg, 2017). In the case of the Antipodean and Gibson’s birds, observed differences in breeding and non-breeding ranges (Walker & Elliott, 2006) combined with strong natal philopatry may reduce opportunities for contact and therefore act as a barrier to gene flow.

Inference of unidirectional gene flow from Adams Island to Disappointment Island indicates the Disappointment Island population was likely founded by the migration of individuals from the main breeding population on Adams Island (Figure 3A; Table SA4). Overall differentiation of the Disappointment Island and Adams Island samples is low compared to the differentiation observed between the Antipodean and Gibson’s subspecies (Figure SA5), and the significance of these results should be interpreted with caution due to a small number of samples collected on Disappointment Island (N = 8). The management implications of the differentiation between the Gibson’s albatross sampling sites are also unclear. However, this finding reveals a previously unidentified avenue for future investigations. Further study should be done to assess whether there are other differences that may be of conservation importance. For example, tracking studies could reveal whether the Disappointment Island birds are utilising similar foraging ranges as the previously tracked Adams Island birds (Walker & Elliott, 2006), and thus whether they are encountering similar threats at sea. This information would also provide further insight into the relationship between at-sea distribution and population structure in these taxa.

### Signatures of selection

Patterns of differentiation at outlier loci and XP-nSL statistics suggest the taxa may be under selection, with several regions of the genome showing signatures of a recent selective sweep. Estimates of absolute genomic divergence were generally low, with low D_XY_ observed across these regions, suggesting that the observed peaks in F_ST_ may at least partially reflect reduced within-population diversity rather than reduced gene flow leading to increased absolute divergence (Cruickshank & Hahn, 2014). However, the observed XP-nSL peaks provide additional evidence for recent selective sweeps in both populations, consistent with divergent selection under different environmental conditions (Ferrer-Admetlla et al., 2014; Szpiech et al., 2021). Although selective sweeps would also be expected to result in reduced π and Tajima’s D estimates, the extremely low genome-wide nucleotide diversity observed in this system may limit the magnitude of these signals and make them difficult to detect. Thus, even in the absence of extensive absolute genomic divergence, incorporating signals from outlier loci and XP-nSL statistics can provide additional insight into evolutionary divergence, particularly adaptive divergence, in populations with extensive shared ancestry that assessment of neutral variation alone may miss.

Putative local adaption may be driven by various environmental or ecological factors. Antipodes Island is at a lower latitude than the Auckland Islands with a higher average temperature and a later, longer period of surrounding ocean productivity that may allow for breeding to continue later in the season. These differences may contribute to the asynchronous timing of breeding between the Auckland Islands and Antipodes Island (Walker & Elliott, 2005) which may also be acting as an extrinsic barrier to gene flow. The notable plumage and morphological differences (Walker et al., unpublished data, 2024) may also indicate traits under selection. Plumage melanism has been associated with adaptive traits in other avian taxa including climate adaptation (Burtt & Ichida, 2004; Delhey, 2019), parasite defence (Briggs et al., 2025; Jacquin et al., 2011), feather strength (Barrowclough & Sibley, 1980; Bonser, 1995) and sexual signalling/mate choice (Cooke et al., 1976; de Zwaan et al., 2019). Similarly, observed differences in wing length may reflect different oceanic wind conditions and foraging range sizes.

Without further study it is not possible to determine what factors are driving this divergence or the traits that may be under selection. However, the candidate genes identified in outlier regions may hold some clues. Several of these candidate genes have been associated with traits in other birds or mammals that are consistent with the observed differences between the taxa (see Supplementary Materials C) including melanism, thermal tolerance, metabolism and morphology. These candidate genes provide biologically plausible hypotheses regarding traits that may underlie the observed genomic differentiation and provide avenues for further research. However, any associations between genes identified in putatively adaptive regions of the genome and these phenotypes remain highly speculative without further study and should be interpreted with caution. Regardless, the evidence of putatively adaptive genomic differentiation presented here is highly relevant for conservation purposes as it highlights the need to preserve genome-wide diversity across the range of Antipodean and Gibson’s albatrosses for long-term resilience and adaptability of populations (Funk et al., 2012).

### Sex chromosome evolution

Sex chromosomes are thought to be important in the evolution of reproductive isolation due to their faster rate of evolution and influence on sexually selected traits (Payseur et al., 2018; Qvarnström & Bailey, 2009). Two of the scaffolds possessing signatures of selection (scaffolds 6 and 22) are putative Z-chromosome scaffolds (Foote et al., 2026), suggesting a potential role for sexual selection in the observed population divergence. Furthermore, patterns of nucleotide diversity and genomic divergence across a region of scaffold 6 present evidence consistent with a recently differentiated evolutionary stratum (a region of recent recombination suppression between the Z- and W-chromosome; Lahn & Page, 1999; Nam & Ellegren, 2008; Schield et al., 2019; Zhou et al., 2014). Interestingly, a recent study in passerines identified an association between recombination suppression of ancestral PARs and the evolution of neo-sex chromosomes (Sigeman et al., 2026) which are increasingly recognised as important drivers of avian evolution (Burley et al., 2023; Gan et al., 2019; Huang et al., 2022; Pala et al., 2012; Sigeman et al., 2020). These findings illustrate how the integration of neutral and putatively adaptive whole-genome data allows us to go beyond merely identifying population structure and enables the formation of hypotheses regarding the mechanisms driving divergence. Future work to resolve the structure of the Z- and W-chromosomes in albatrosses would enable the identification of PARs, evolutionary strata and structural variation such as neo-sex chromosomes, providing further insight into the role of sex chromosome evolution and its function in population divergence and adaptation in albatrosses and other Procellariiformes.

### Demographic trends

Evaluation of contemporary N_e_ estimates, as well as recent population history revealed that the Antipodean subspecies has comprised a larger effective population than the Gibson’s subspecies over time (Figure 3B). This is somewhat surprising as it contradicts census estimates showing a larger number of Gibson’s breeding pairs (Elliott et al., 2025; Rexer- Huber et al., 2025; Richard, 2021). Such a contrast may be explained by differences in key demographic rates. Since the populations were first monitored in the early 1990s, average annual nesting success has been higher in the Antipodean subspecies (Parker et al., 2023; Walker et al., 2023). Although the Antipodean subspecies appears to be suffering more severe ongoing declines, this is largely attributed to high female mortality in fisheries bycatch and prior to the observed population size crash in 2005 this population was increasing at a much faster rate than the Gibson’s.

The observed demographic trends in combination with our findings indicate that census estimates may not accurately represent current or past population productivity and also show that the taxa are demographically independent. Ovenden (2013) demonstrated that when the number of migrants is very low, genetics provides good insight into the demographic connectivity between populations. Only a small number of migrants per generation are required to genetically connect populations and homogenise gene pools (Lowe & Allendorf, 2010; Wright, 1949). Thus, the observed genetic differentiation and inferred lack of gene flow, alongside the varying demographic rates demonstrated over the long-term population study, suggest the subspecies are more distinct than previously thought.

### Conservation implications

Information on population structure and discrete breeding units is essential for appropriate conservation management of threatened populations (Funk et al., 2012; Morid, 1994). In the past, the taxonomic status of the Antipodean and Gibson’s albatross was reclassified from species to subspecies on the basis that although significant genetic differentiation was identified, the differences were not fixed. The recommendation was made that the populations be treated as distinct management units in recognition of the significant allele frequency differences and differences in at-sea distribution, likely requiring different management approaches (Burg & Croxall, 2004). In NZ, the subspecies are managed separately, with efforts to address the threats at a higher taxonomic resolution (Richard et al., 2024; Robertson et al., 2021). However, one of the biggest threats these taxa face is overlap with pelagic longline fisheries outside the jurisdiction of the NZ EEZ (Bose & Debski, 2020; Walker & Elliott, 2006) where conservation action remains primarily focussed at the species level (IUCN, 2012, 2022; UNEP/CMS, 2020) and population declines are continuing at an alarming rate (Parker et al., 2023; Walker et al., 2023).

This study confirms the significant genetic differentiation identified by Burg and Croxall (2004), but helps to further resolve the population structure by demonstrating population differentiation with a lack of inferred gene flow, as well as potential adaptive divergence. The threats to the populations are well described (Bose & Debski, 2022; Parker et al., 2023; Walker et al., 2023). Thus, the combined neutral and putatively adaptive genomic data presented in this study are important because they show that not only are the populations genomically differentiated and have been demographically independent for thousands of years, but they are also on different evolutionary trajectories and may not be receiving adequate protection while they are recognised as a single Evolutionarily Significant Unit (ESU; Funk et al., 2012). Evidence of genomic divergence agrees with other lines of evidence suggesting differentiation between the taxa which, taken together, may warrant an integrative taxonomic reassessment (Walker et al., unpublished data, 2024; Walker & Elliott, 2005, 2006). Should a reassessment lead to a taxonomic revision, this will have implications for their conservation management as an updated threat status for each new species may reveal more urgent conservation action is required to protect them from threats.

Genetic diversity is an important measure of the genetic health of a population, and population declines can lead to a loss of genetic diversity and subsequent reduction in fitness (Charlesworth & Charlesworth, 1987; Reed & Frankham, 2003; van der Valk et al., 2019). Although low overall genomic diversity was identified, no concerning reduction in heterozygosity leading to inbreeding was observed in either taxon in the current whole-genome dataset. However, it is not possible to conclude from these data that the most recent declines have not resulted in loss of diversity, as loss of alleles and reduction in heterozygosity occurs over successive generations following population bottlenecks (Nei et al., 1975). The most concerning declines have occurred only over the past 20 years (Parker et al., 2023; Walker et al., 2023), and due to the long generation times of albatrosses (approx. 25 years; Bird et al., 2020), it is likely that the genetic effects of these declines will not be reflected in the current genomic dataset. This dataset is valuable as it provides a benchmark for current genetic diversity, against which potential future losses in diversity can be monitored. It is also important to note that genomics is only one tool and is most valuable for conservation management when applied alongside other monitoring and tracking efforts.

### Conclusions

Our genomic dataset revealed significant genetic differentiation between the Antipodean and Gibson’s albatross subspecies at both neutral and putatively adaptive (outlier) loci. A lack of inferred gene flow indicates that the low levels of genomic differentiation observed may be due to recent shared ancestry and historically low levels of genetic variation, rather than ongoing gene flow and suggest the subspecies are not well connected genetically and are demographically independent. Advances in whole-genome sequencing have revealed similar observations across other seabirds, including other albatrosses, which provides further insight into the evolution and drivers of population structure in Southern Ocean seabirds. This study provides the first high-quality whole-genome dataset for assessing population structure in *Diomedea* albatrosses, allowing high confidence in the identified patterns of structure and setting a benchmark for assessment of other *Diomedea* species. This genomic dataset also allowed integration of putatively adaptive (outlier) alongside genome-wide neutral markers for the Antipodean and Gibson’s albatross, providing a more complete picture of genomic and evolutionary divergence than either approach alone. More broadly, this study demonstrates how the complementary evidence of population structure, genetic connectivity and potential divergent selection can help to reveal unique genetic variation and evolutionary processes, providing a powerful framework for resolving recently diverged taxa beyond what can be achieved through assessment of neutral variation alone.

## Acknowledgements

Sampling for this research was undertaken under an agreement between Te Rūnanga o Ngāi Tahu and the Department of Conservation NZ. We thank Ngāi Tahu for their support of this research. We also thank the following people and organizations for their support with this work; Igor Debski and Johannes Fischer (NZ Department of Conservation), Genomics Aotearoa, Dini Senanayake and Vicky Fan (New Zealand eScience Infrastructure) and BirdsNZ. This work was supported by the NZ Department of Conservation as part of the Conservation Services Programme project INT2019/02. Geoff Chambers thanks VUW for alumnus scholar support.

## Data Accessibility

Toroa (a range of albatross species, including Antipodean and Gibson’s albatross) are taonga (treasured) species for Ngāi Tahu (the Indigenous Peoples of southern Aotearoa New Zealand). The genomes represent the whakapapa (genealogy) of these taonga species and so are considered taonga themselves. Rangatiratanga (sovereignty, chieftainship, and/or self- determination) and kaitiakitanga (guardianship, practices informed by centuries of experiences to achieve intergenerational sustainability) over these taonga remain with Ngāi Tahu. Accordingly, tikanga Māori (customs/ethical system of common law and practice/guidance on practicing kaitiakitanga) determines how people interact with them. To ensure that management and storage of these samples and any subsequently generated genomic data follow tikanga Māori, including to ensure that rangatiratanga remains with Ngāi Tahu, a formal sample and data management agreement has been signed by Te Rūnanga o Ngāi Tahu and the New Zealand Department of Conservation. This agreement outlines that the genomes will be stored at a culturally informed facility (Aotearoa Genomic Data Repository; https://data.agdr.org.nz/), where the data can be made available at the discretion of Ngāi Tahu upon request.

Accordingly, we have deposited the primary data underlying these analyses as follows:

- Raw reads and all associated metadata: Aotearoa Genomic Data Repository (AGDR) under project AGDR00069).
- All scripts associated with bioinformatic analyses: Zenodo https://doi.org/10.5281/zenodo.22251389

## Author contributions

Imogen Foote (Research design, Data analysis, Project administration, Writing – original draft, Writing – review & editing), Tom Oosting (Data analysis, Writing – review & editing), Geoff Chambers (Research design, Supervision, Writing – review & editing), Kath Walker (Sample collection, Writing – review & editing), Graeme Elliott (Sample collection, Writing – review & editing), Kalinka Rexer-Huber (Sample collection, Writing – review & editing), Graham Parker (Sample collection, Writing – review & editing), Peter Ritchie (Research design, Funding acquisition, Project administration, Supervision, Writing – review & editing).

## Ethics approval statement

Sampling methods were approved by the Victoria University of Wellington Animal Ethics Committee (VUW AEC 30055).

