## Supplementary Materials A for "Integrating genome-wide neutral and putatively adaptive variation\ resolves population structure and differentiation in a recently diverged albatross complex"

**Table of Contents:**

| **Figure SA1.** Per site quality metrics of all polymorphic SNPs (7,285,599 SNPs; before quality filtering) showing **A)** Site quality (Phred Score), **B)** Site missingness and **C)** Mean depth per site | Page 3 |
| --- | --- |
| **Figure SA2.** Impact of different quality filtering thresholds on observed population structure of the Antipodean and Gibson’s albatross. Samples are grouped by sampling island. **A)** Missingness 0.99, MAF 0.05, Q30, N SNPs = 2,224,100, **B)** Missingness 0.95, MAF 0.05, Q30, N SNPs = 2,400,435, **C)** Missingness 0.95, MAF 0.01, Q30 N SNPs = 4,300,448 and **D)** Missingness 0.90, MAF 0.01, Q30, N SNPs = 4,312,932. | Page 4-5 |
| **Figure SA3.** Mean sequence quality (Phred) scores from all individuals assessed using FastQC and summarised and visualised with MultiQC. Quality assessment was performed prior to adapter trimming. | Page 6 |
| **Figure SA4.** Quality statistics per individual of all polymorphic sites (7,285,599 SNPs; before filtering) showing **A)** mean sequencing depth, **B)** inbreeding coefficient, **C)** observed heterozygosity and **D)** proportion of missing genotypes. Note: depth statistic presented from calculations after high-depth outliers were randomly downsampled. | Page 7 |
| **Figure SA5.** Pairwise weighted F_ST_ estimates between **A)** all sampling sites and **B)** sampling islands of Antipodean and Gibson’s albatross | Page 8 |
| **Figure SA6.** Patterns of D_XY_, F_ST_ and nucleotide diversity (π) across scaffold 6 (putative Z-chromosome scaffold) in **A)** male and **B)** female Antipodean and Gibson’s albatrosses. | Page 9 |
| **Figure SA7. A)** Heterozygosity and **B)** mean sequencing depth across scaffold 6 (putative Z-chromosome scaffold) in male and female Antipodean and Gibson’s albatrosses. | Page 10 |
| **Table SA1.** Analysis of Molecular Variance (AMOVA) between Antipodean and Gibson's albatross populations using 381,176 neutral SNPs. | Page 11 |
| **Table SA2.** Analysis of Molecular Variance (AMOVA) between Antipodean and Gibson's albatross populations using 57 outlier SNPs. | Page 11 |
| **Table SA3.** Cross-validation error from ADMIXTURE analyses. Cross-validation error was performed for K = 1-5. Lowest cross-validation error (shown in bold) represents the best predictive value of K (number of ancestral populations) for the dataset. | Page 12 |
| **Table SA4.** Inferred mean migration rates with approximate 95% credible intervals estimated between subspecies (Antipodean and Gibson’s albatross) and islands (Antipodeans = Antipodes, Gibson’s = Adams, Disappointment) using BA3-SNPs. Approximate credible intervals were calculated as mean±1.96×SD per the BayeAss Edition 3.0 User’s Manual (Rannala, 2007). Values in bold show significant estimated migration (95% credible intervals do not include zero). Proportion of non-migrants for each population shown along diagonal (light shading). | Page 12 |
| **Table SA5.** Contemporary effective population size (N_e_) of Antipodean and Gibson's albatross calculated in NeEstimator2. Upper and lower confidence intervals are based on jacknife resampling. | Page 13 |

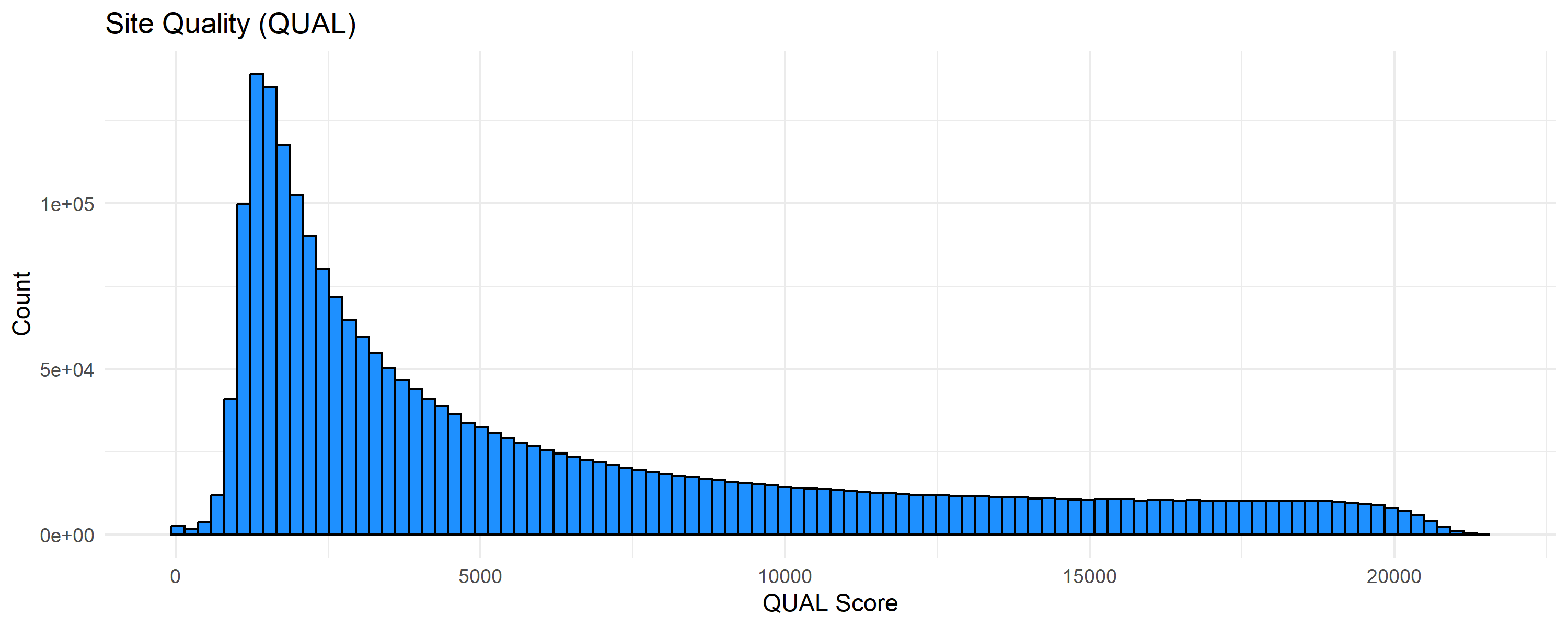

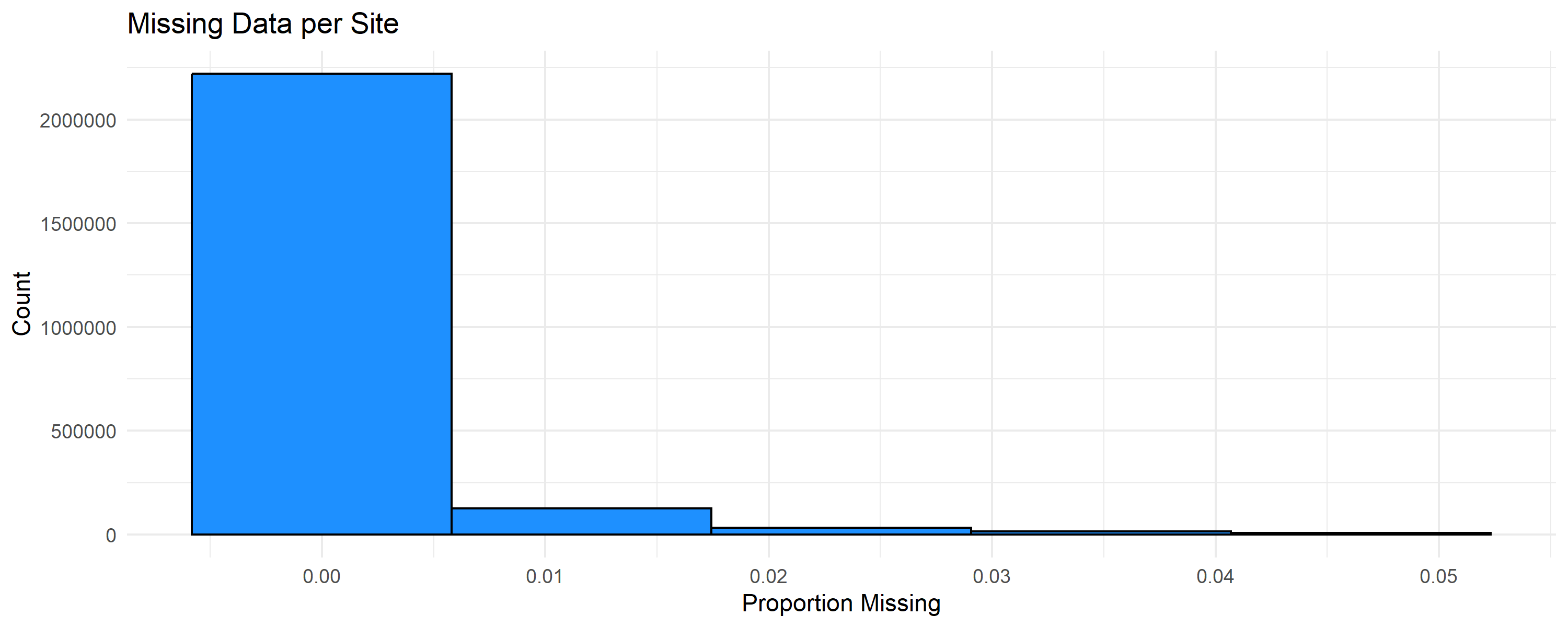

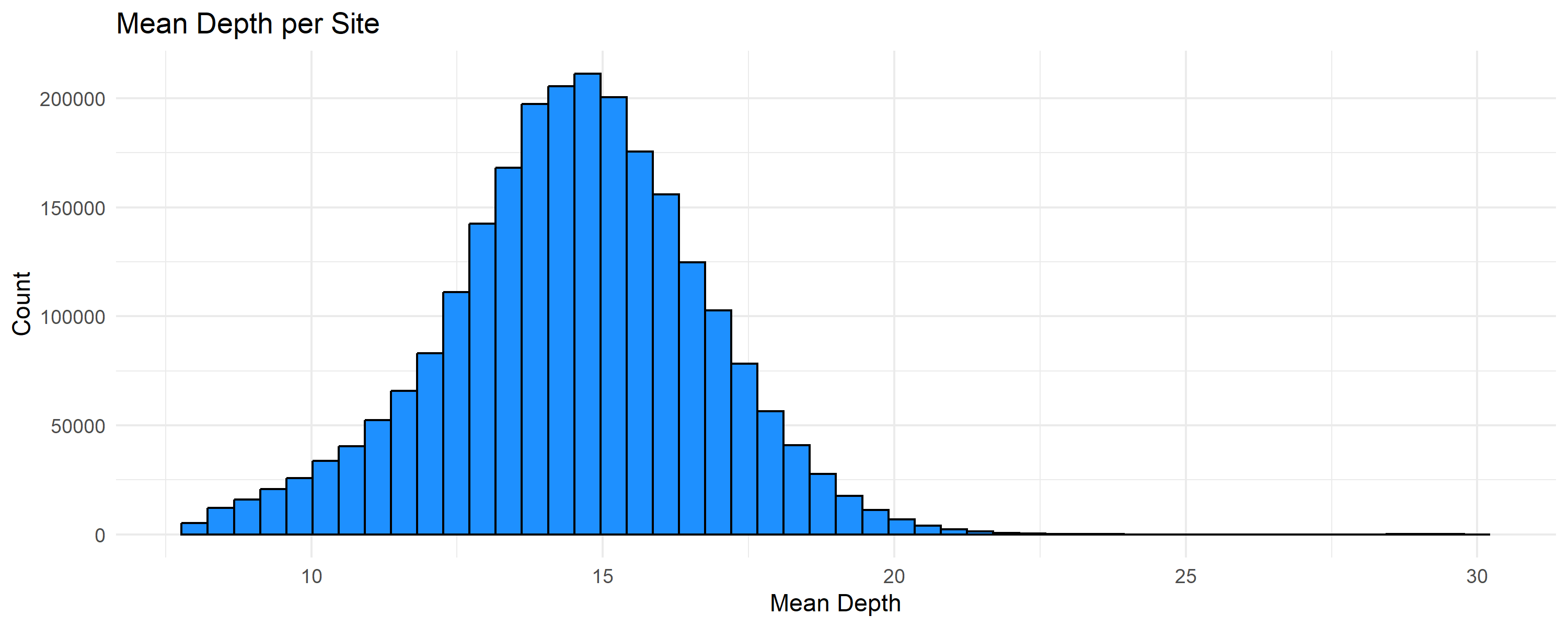

**A)**

**B)**

**C)**

**Figure SA1.** Per site quality metrics of all polymorphic SNPs (7,285,599 SNPs; before quality filtering) showing **A)** Site quality (Phred Score), **B)** Site missingness and **C)** Mean depth per site.

**A).** Missingness 0.99, MAF 0.05, Q30

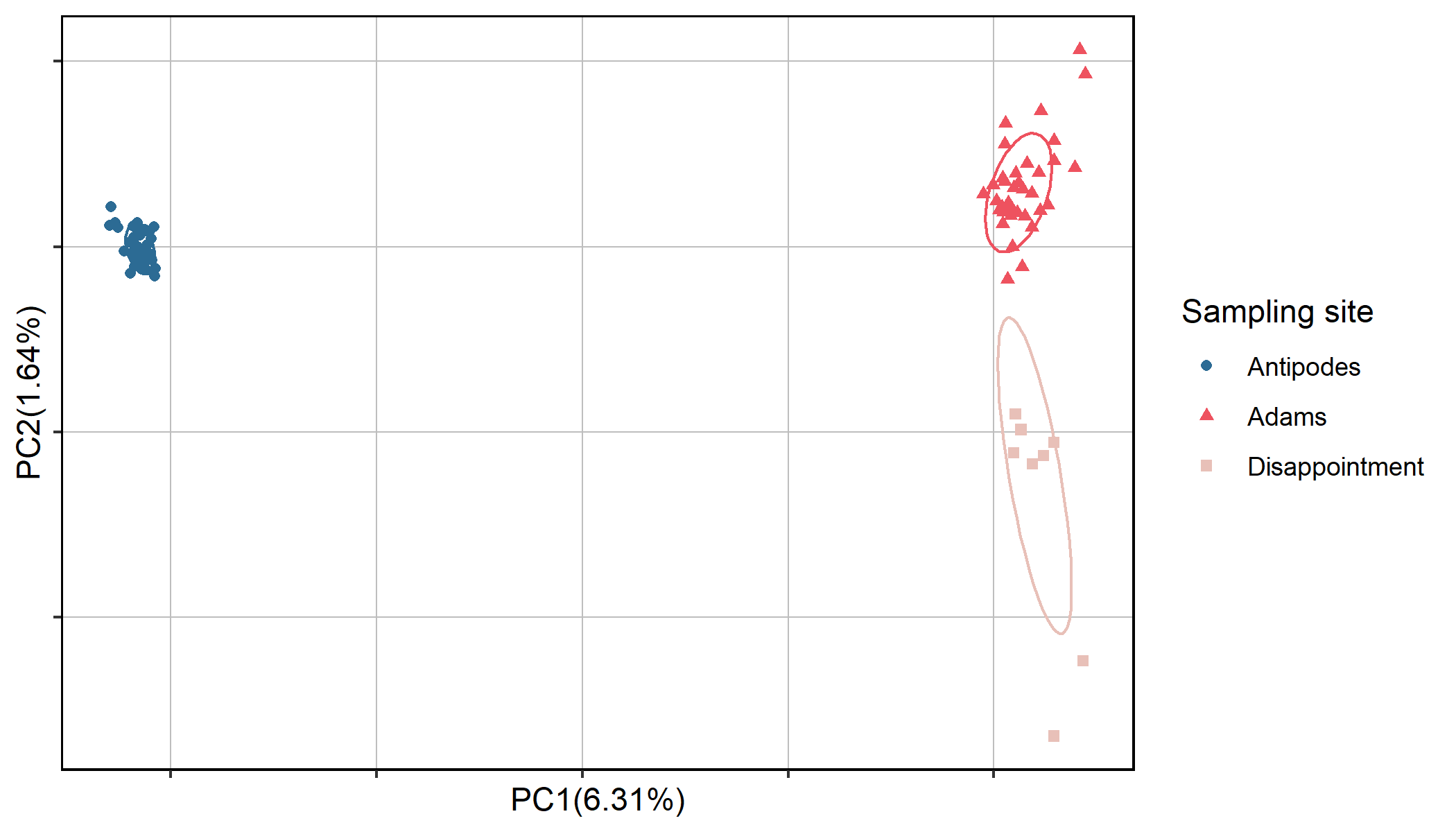

**B)** Missingness 0.95, MAF 0.05, Q30

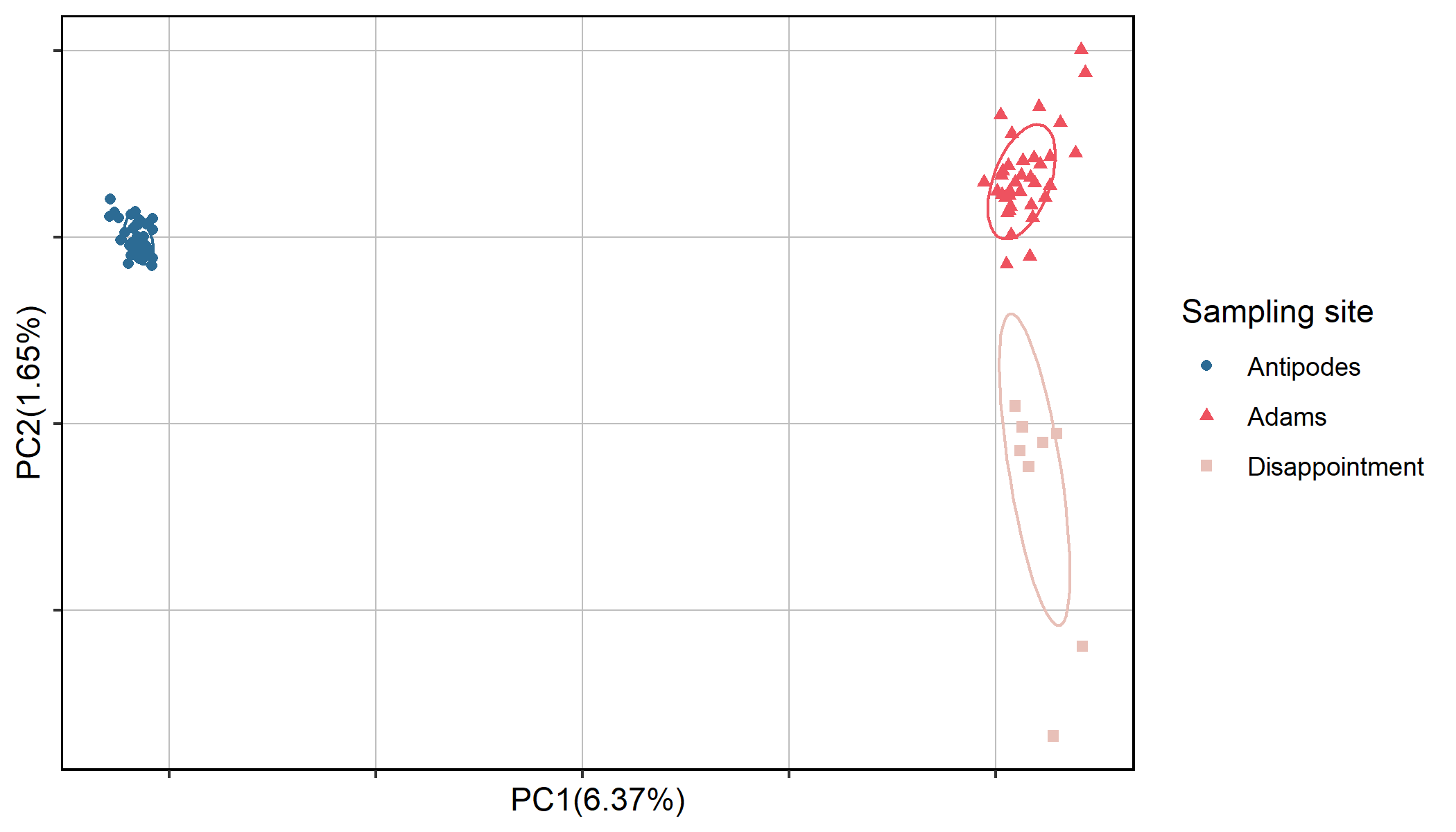

**C)** Missingness 0.95, MAF 0.01, Q30

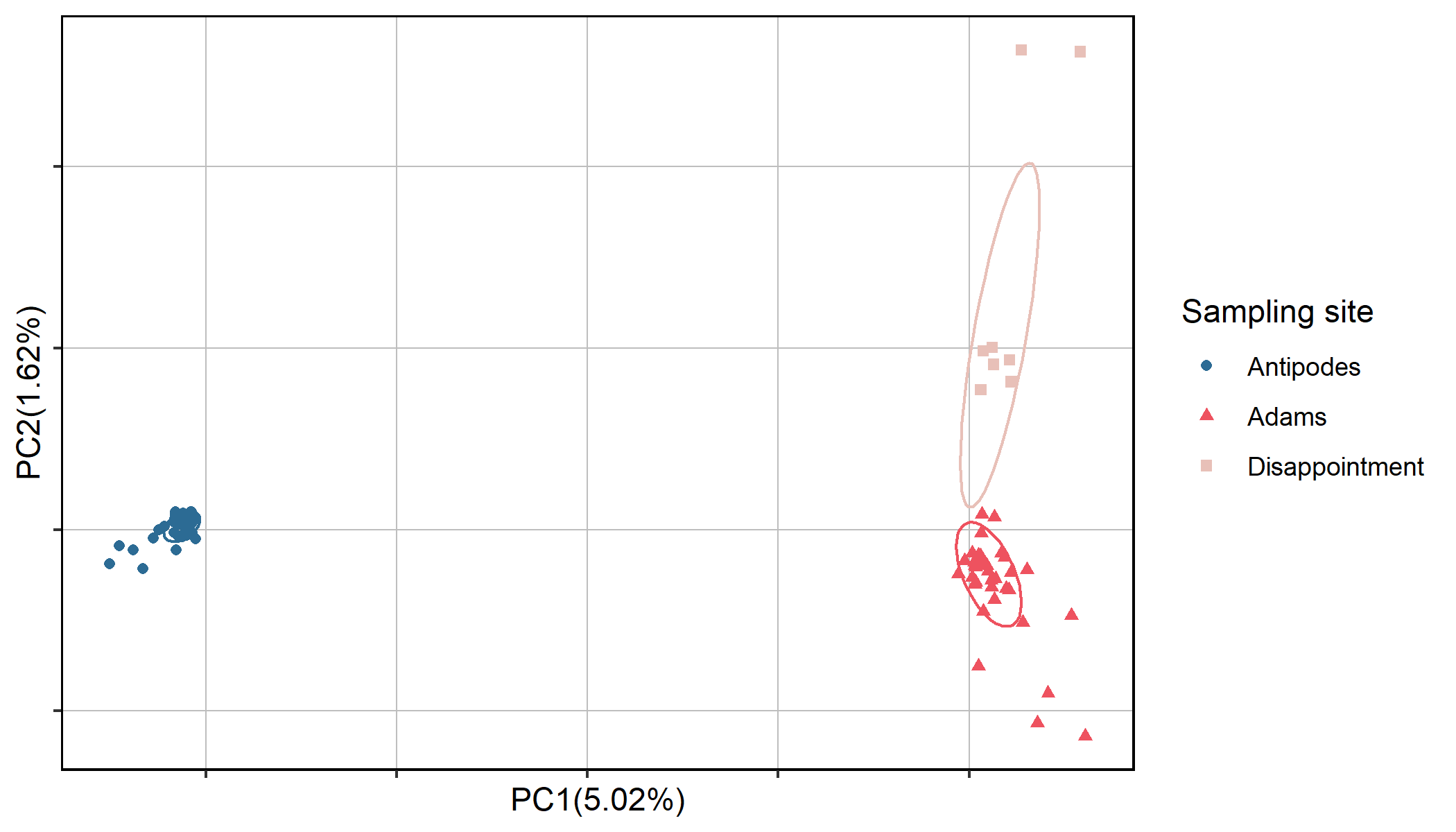

**D)** Missingness 0.90, MAF 0.01, Q30

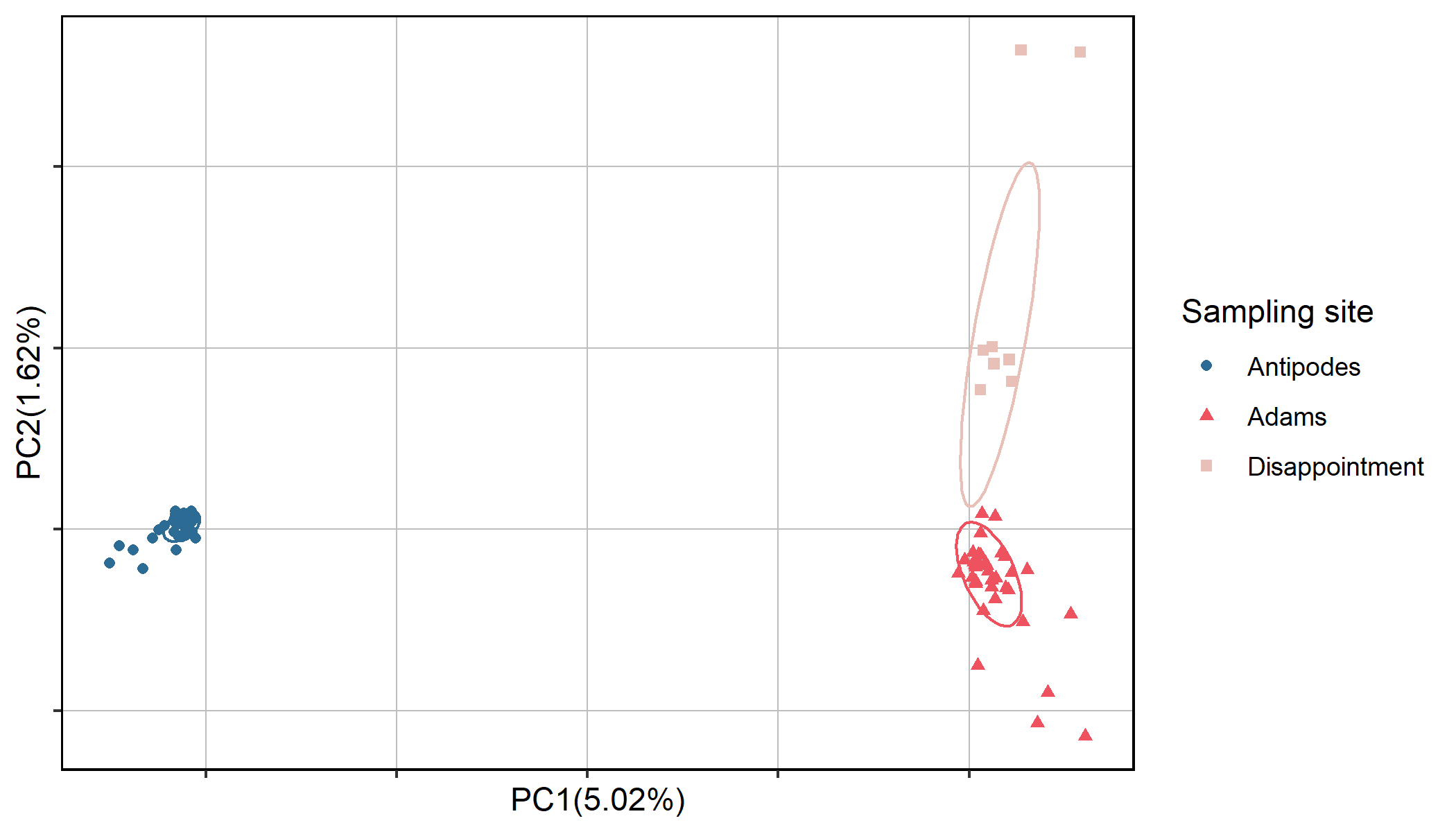

**Figure SA2.** Impact of different quality filtering thresholds on observed population structure of the Antipodean and Gibson’s albatross. Samples are grouped by sampling island. **A)** Missingness 0.99, MAF 0.05, Q30, N SNPs = 2,224,100, **B)** Missingness 0.95, MAF 0.05, Q30, N SNPs = 2,400,435, **C)** Missingness 0.95, MAF 0.01, Q30 N SNPs = 4,300,448 and **D)** Missingness 0.90, MAF 0.01, Q30, N SNPs = 4,312,932.

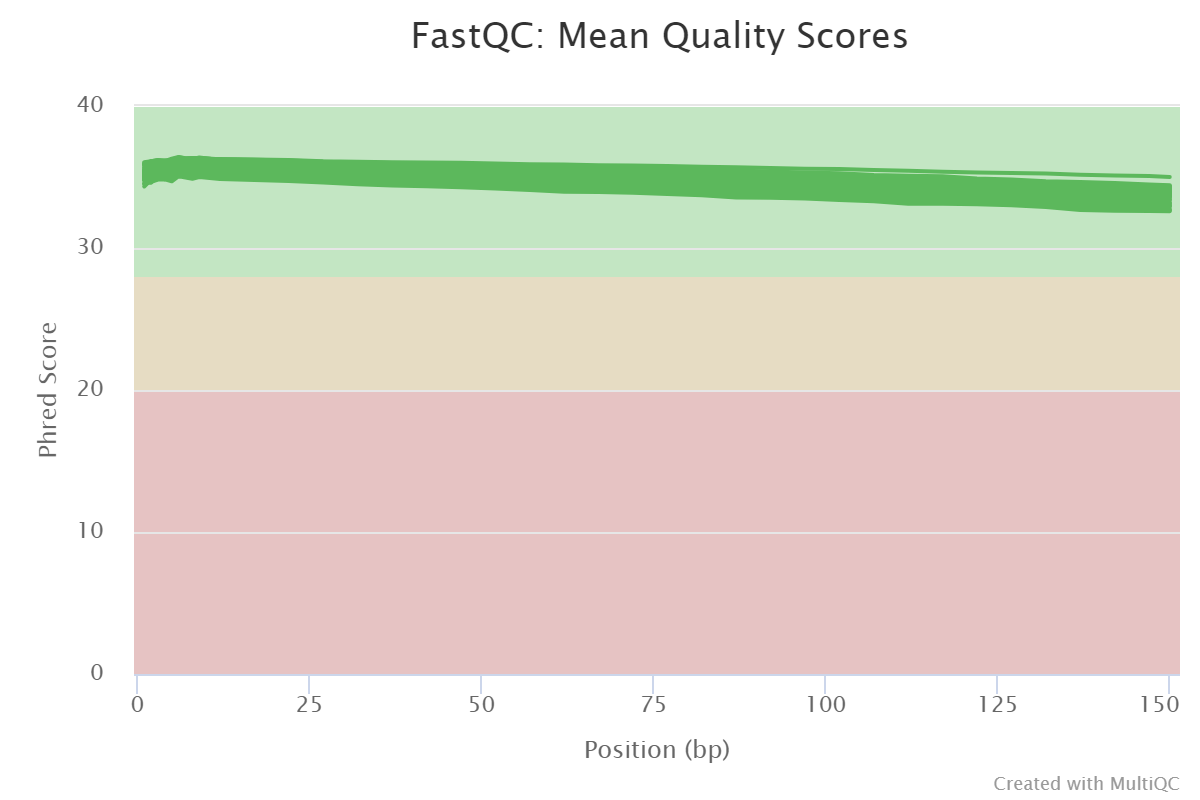

**Figure SA3**. Mean sequence quality (Phred) scores from all individuals assessed using FastQC and summarised and visualised with MultiQC. Quality assessment was performed prior to adapter trimming.

**A)**

**B)**

**C)**

**D)**

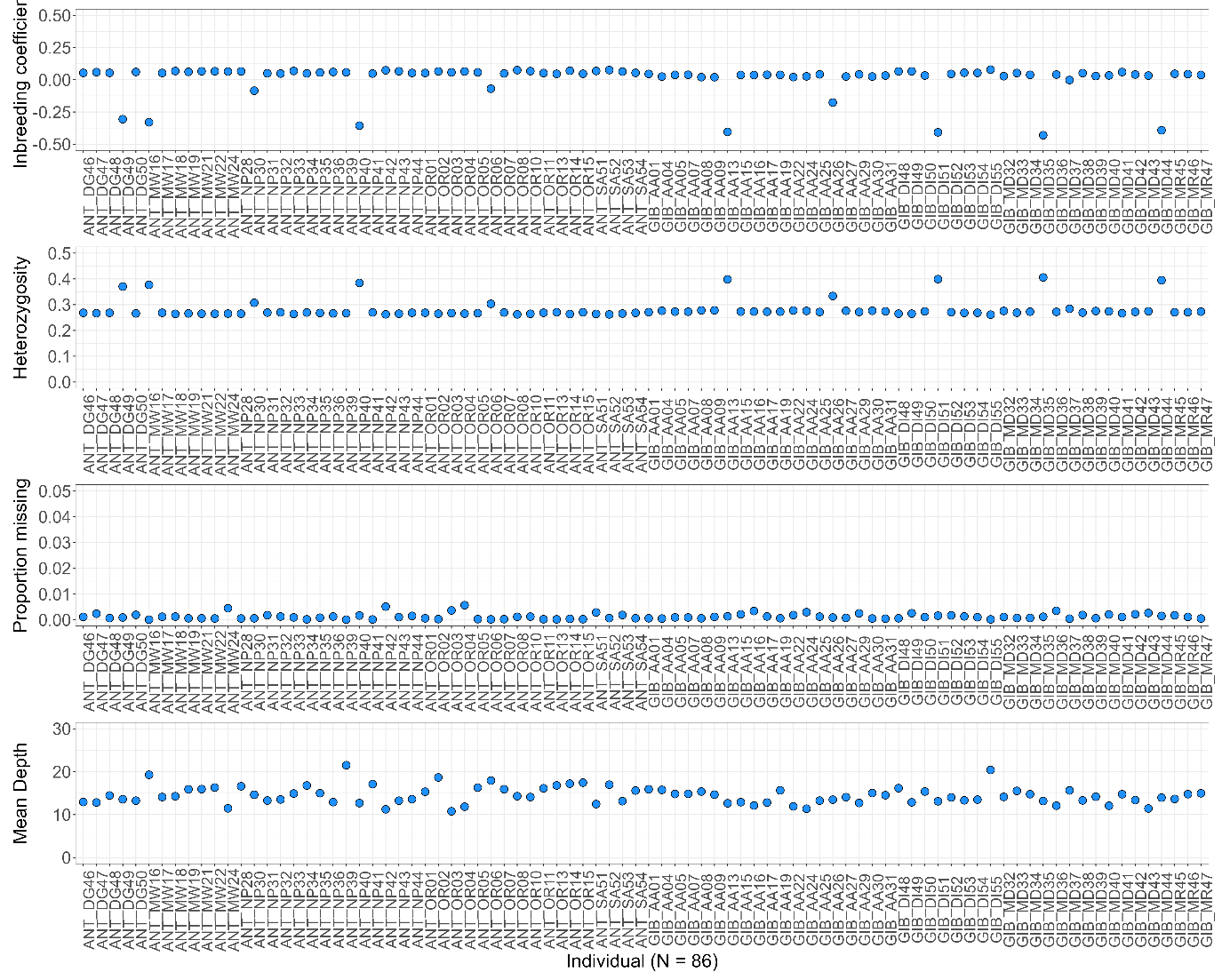

**Figure SA4.** Quality statistics per individual of all polymorphic sites (7,285,599 SNPs; before filtering) showing **A)** mean sequencing depth, **B)** inbreeding coefficient, **C)** observed heterozygosity and **D)** proportion of missing genotypes. Note: depth statistic presented from calculations after high-depth outliers were randomly downsampled.

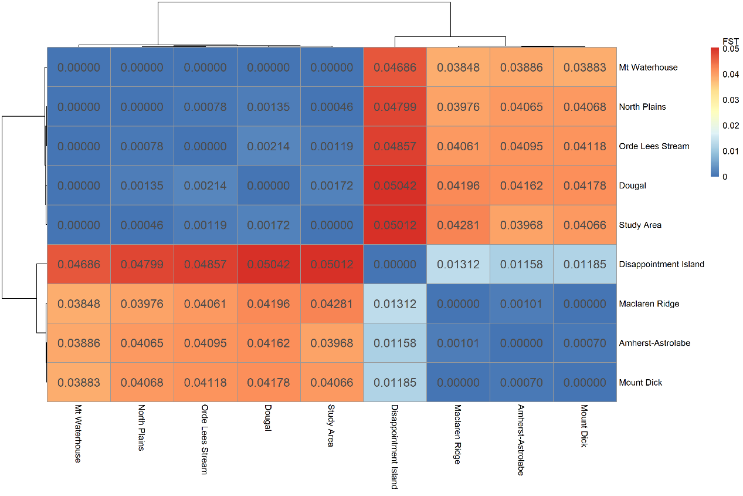

**Figure SA5.** Pairwise weighted F_ST_ estimates between **A)** all sampling sites and **B)** sampling islands of Antipodean and Gibson’s albatross

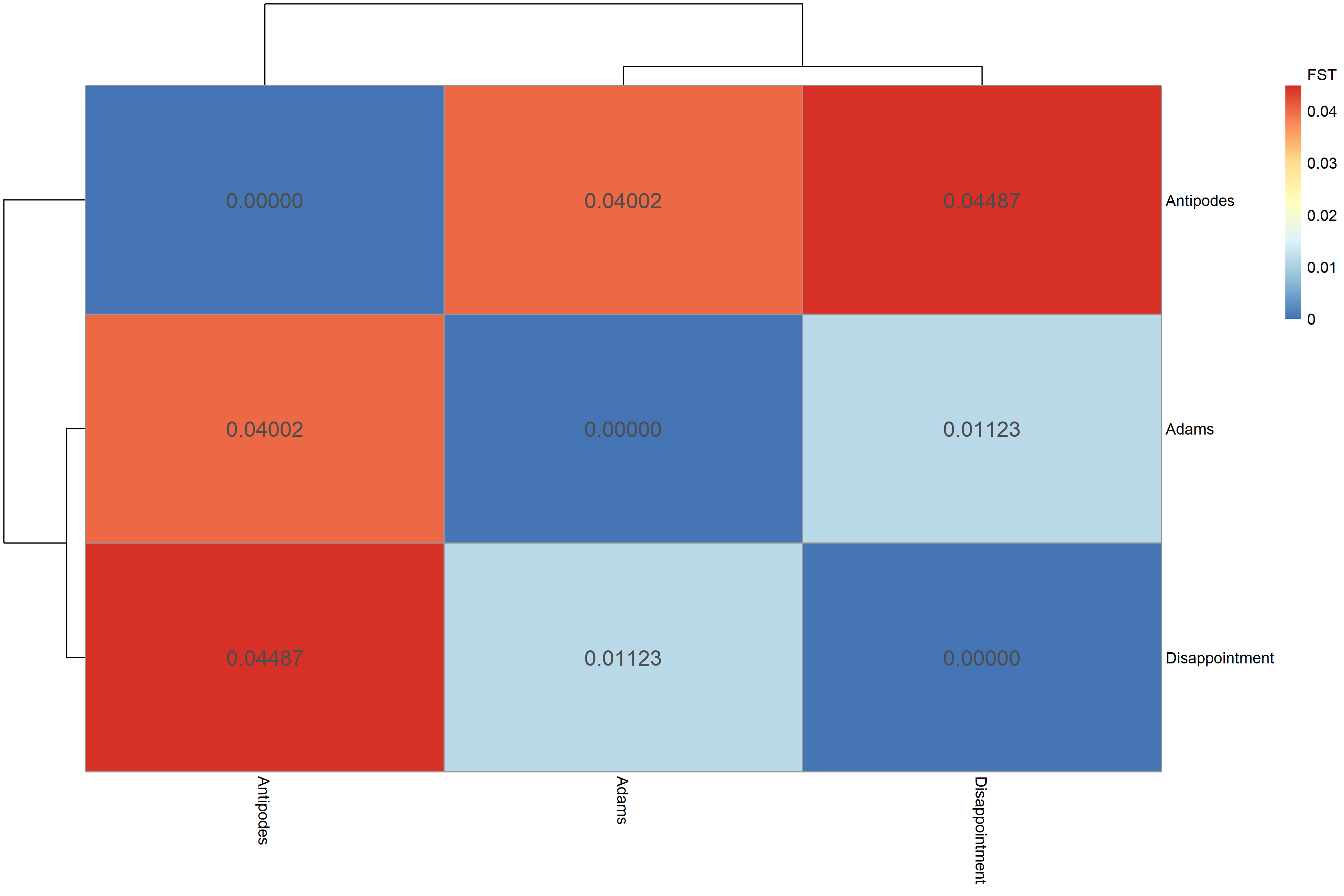

**A)**

**B)**

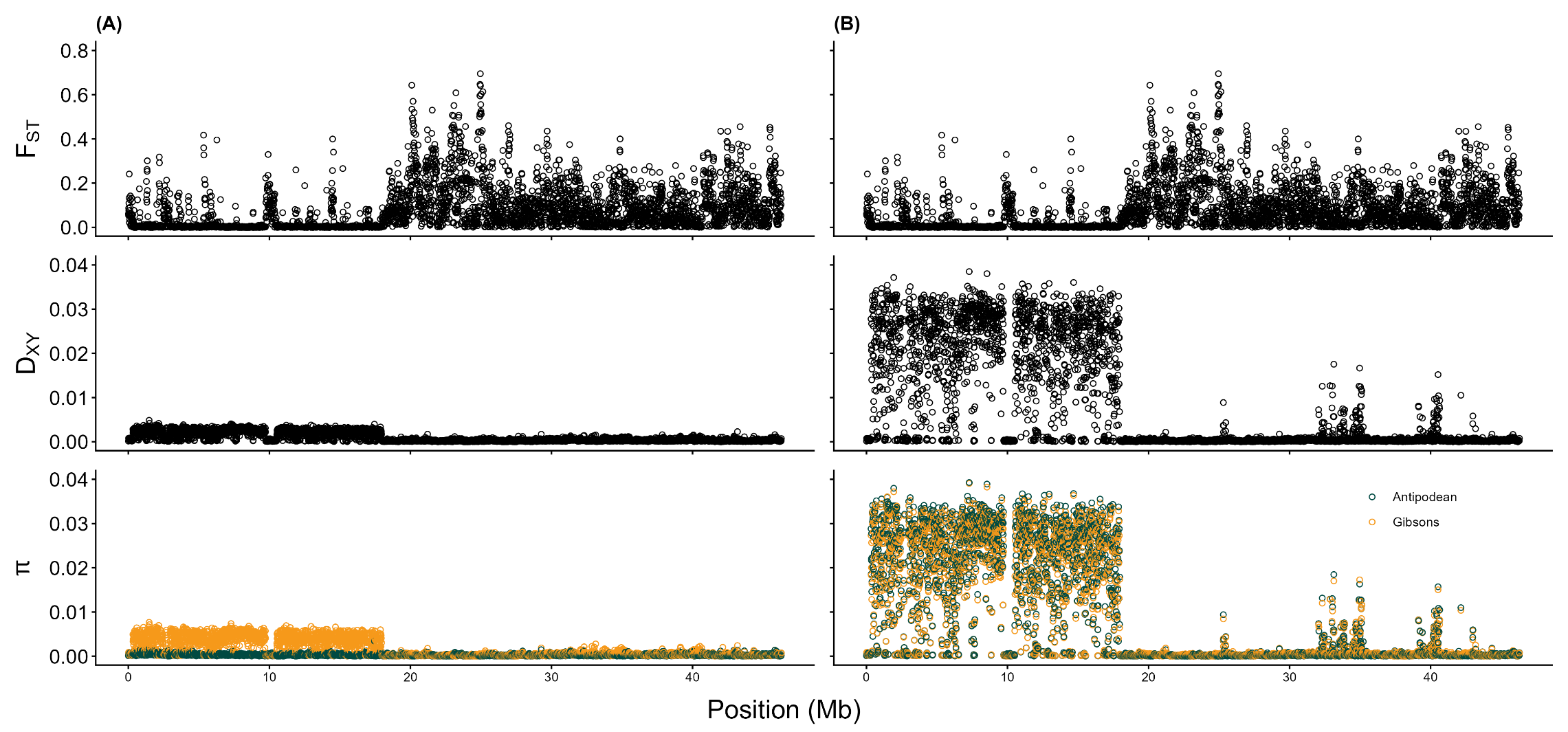

**Figure SA6.** Patterns of D_XY_, F_ST_ and nucleotide diversity (π) across scaffold 6 (putative Z-chromosome scaffold) in **A)** male and **B)** female Antipodean and Gibson’s albatrosses.

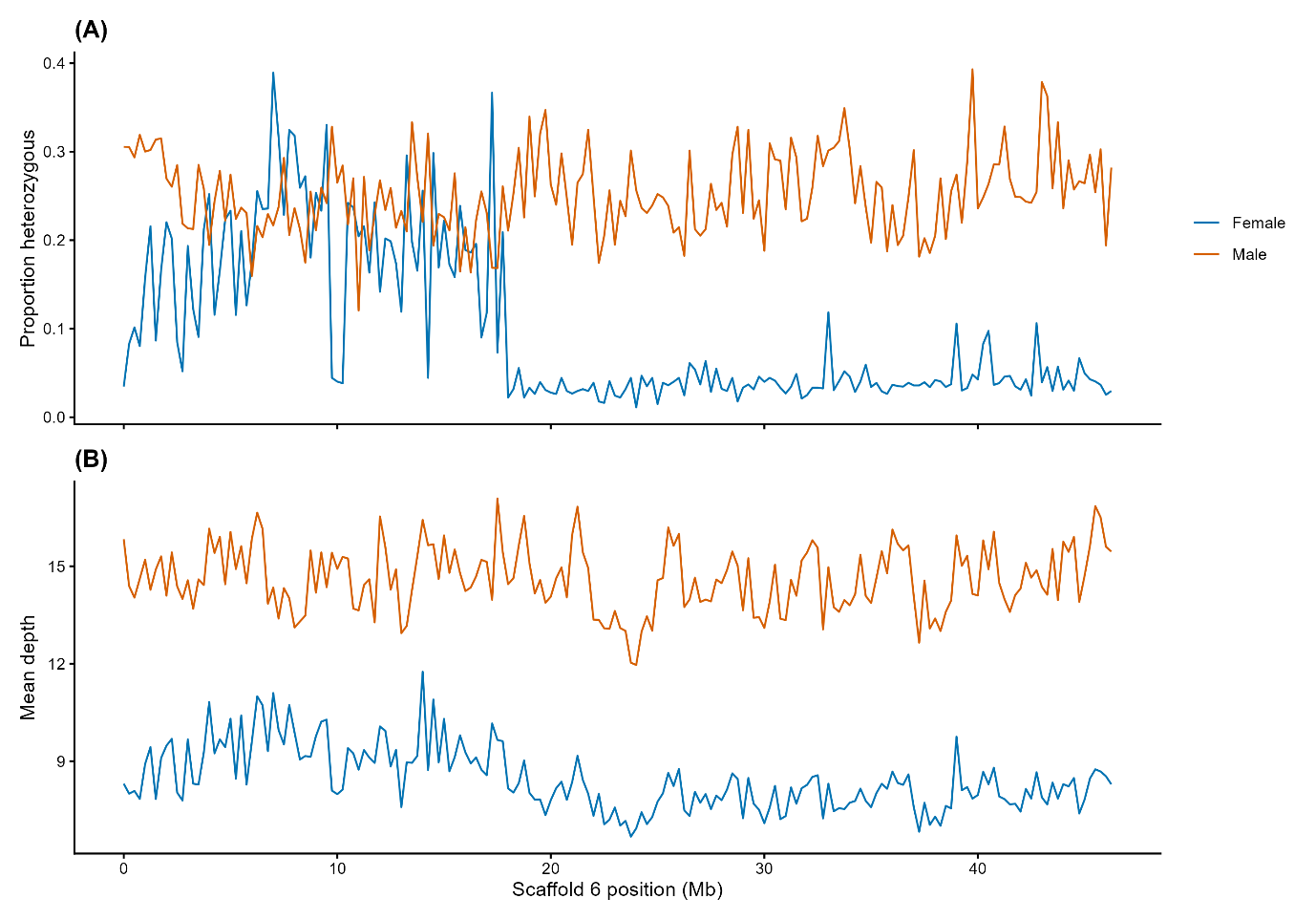

**Figure SA7.** **A)** Heterozygosity and **B)** mean sequencing depth across scaffold 6 (putative Z-chromosome scaffold) in male and female Antipodean and Gibson’s albatrosses.

**Table SA1.** Analysis of Molecular Variance (AMOVA) between Antipodean and Gibson's albatross populations using 381,176 neutral SNPs.

| Source of variation | d.f. | Sum of squares | Mean sum of squares | Variance components* | Percentage of variation | Φ | *p* |
| --- | --- | --- | --- | --- | --- | --- | --- |
| Between populations | 1 | 262,393.3 | 262,39.3 | 2,402.8 | 4.00 | 0.0098 | 0.001 |
| Between samples within populations | 84 | 4,682,869.8 | 55,748.5 | -1,817.5 | -3.03 | -0.0316 | 0.929 |
| Within samples | 86 | 5,106,982.9 | 59,383.5 | 59,383.5 | 99.02 | 0.0400 | 0.660 |
| **Total** | **171** | **10,052,246.0** | **58,785.1** | **59,968.8** | **100.00** |  |  |
| d.f. = degrees of freedom, *slightly negative covariance can occur in the absence of genetic structure, Φ = phi statistic, *p* = p-value. | | | | | | | |

**Table SA2.** Analysis of Molecular Variance (AMOVA) between Antipodean and Gibson's albatross populations using 57 outlier SNPs.

| Source of variation | d.f. | Sum of squares | Mean sum of squares | Variance components* | Percentage of variation | Φ | *p* |
| --- | --- | --- | --- | --- | --- | --- | --- |
| Between populations | 1 | 743.2 | 743.2 | 8.54 | 50.70 | 0.5092 | 0.001 |
| Between samples within populations | 84 | 700.7 | 8.3 | 0.04 | 0.21 | 0.0042 | 0.421 |
| Within samples | 86 | 711.3 | 8.3 | 8.27 | 49.08 | 0.5071 | 0.001 |
| **Total** | **171** | 2,155.2 | **12.6** | **16.85** | **100.00** |  |  |
| d.f. = degrees of freedom, *slightly negative covariance can occur in the absence of genetic structure, Φ = phi statistic, *p* = p-value. | | | | | | | |

**Table SA3.** Cross-validation error from ADMIXTURE analyses. Cross-validation error was performed for K = 1-5. Lowest cross-validation error (shown in bold) represents the best predictive value of K (number of ancestral populations) for the dataset.

| K | CV error |
| --- | --- |
| 1 | 0.52440 |
| **2** | **0.51624** |
| 3 | 0.54971 |
| 4 | 0.57667 |
| 5 | 0.61184 |

**Table SA4.** Inferred mean migration rates with approximate 95% credible intervals estimated between subspecies (Antipodean and Gibson’s albatross) and islands (Antipodeans = Antipodes, Gibson’s = Adams, Disappointment) using BA3-SNPs. Approximate credible intervals were calculated as mean±1.96×SD per the *BayeAss Edition 3.0 User’s Manual* (Rannala, 2007). Values in bold show significant estimated migration (95% credible intervals do not include zero). Proportion of non-migrants for each population shown along diagonal (light shading).

|  | *To* | Antipodean | Gibson’s | Antipodes | Adams | Dis. |
| --- | --- | --- | --- | --- | --- | --- |
| *From* | Antipodean | 0.9926 (0.9785 – 1.0067) | 0.0076  (-0.0069 – 0.0221) |  |  |  |
|  | Gibson’s | 0.0074  (-0.0067 – 0.0215) | 0.9924 (0.9779 – 1.0069) |  |  |  |
|  | Antipodes |  |  | 0.9855 (0.9679 – 1.0031) | 0.0088  (-0.0081 – 0.0257) | 0.0302  (-0.0239 – 0.0843) |
|  | Adams |  |  | 0.0072  (-0.0067 – 0.0211) | 0.9824 (0.9589 – 1.0059) | **0.2727 (0.2000 – 0.3454)** |
|  | Disappointment |  |  | 0.0073  (-0.0066 – 0.0212) | 0.0088  (-0.0081 – 0.0257) | 0.6970 (0.6427 – 0.7513) |

**Table SA5.** Contemporary effective population size (N_e_) of Antipodean and Gibson's albatross calculated in NeEstimator2. Upper and lower confidence intervals are based on jackknife resampling.

| Population | N_e_ | CI for N_e_ | |
| --- | --- | --- | --- |
|  |  | Lower | Upper |
| Antipodean | 2,258 | 1,043 | ∞ |
| Gibson’s | 1,157 | 598 | 11,669 |

**Separate Supplementary Document Files**

**Supplementary Materials B.** Population genomic analysis results when reads are mapped to the Gibson’s albatross reference genome.

**Supplementary Materials C.** An Excel spreadsheet with the names, genomic positions and putative function of genes identified within 10,000 bp windows showing extreme XP-nSL values and outlier SNPs identified by Baypass and pcadapt.
