## Supplementary Materials B for "Integrating genome-wide neutral and putatively adaptive variation\ resolves population structure and differentiation in a recently diverged albatross complex"

**Table of Contents:**

| **Figure SB1.** Principal Component Analysis (PCA) based on 524,726 neutral SNPs showing the genetic clustering of Antipodean albatross (left cluster) and Gibson’s albatross (right cluster) with reads aligned to Gibson’s albatross reference genome. Samples are grouped by sampling island. | Page 3 |
| --- | --- |
| **Figure SB2.** ADMIXTURE plot for K = 2 showing the ancestry proportions of individuals from Antipodean and Gibson’s albatross populations using 524,726 neutral SNPs with reads aligned to Gibson’s albatross reference genome. Solid black lines indicate groupings of individuals from each sampling site. DG = Dougal, MW = Mt Waterhouse, NP = North Plains, OR = Orde Lees Stream, SA = Study Area, AA = Amherst Amsterdam, DI = Disappointment Island, MD = Mt Dick, MR = Maclaren Ridge | Page 3 |
| **Figure SB3.** Pairwise weighted FST estimates between **A)** all sampling sites and **B)** sampling islands of Antipodean and Gibson’s albatross with reads aligned to Gibson’s albatross reference genome. | Page 4 |
| **Figure SB4.** Principal Components Analysis (PCA) based on 99 outlier SNPs showing the genetic clustering of Antipodean albatross (left cluster) and Gibson’s albatross (right cluster) with reads aligned to Gibson’s albatross reference genome. Samples are grouped by sampling island. | Page 5 |
| **Figure SB5.** Principal Components Analysis (PCA) based on 99 outlier SNPs showing the genetic clustering of Antipodean albatross (left cluster) and Gibson’s albatross (right cluster) with reads aligned to Gibson’s albatross reference genome. Samples are grouped by sampling island. | Page 5 |
| **Table SB1.** Genetic diversity statistics calculated using a dataset of 524,726 neutral SNPs or a genome-wide SNP dataset (including invariant sites) for the Antipodean and Gibson’s albatross with reads aligned to Gibson’s albatross reference genome.  H_o_ = observed heterozysity (SE), H_e_ = expected heterozygosity (SE), π = average nucleotide diversity, θ = average Watterson’s estimator, F_IS_ = inbreeding coefficient (1 − H_o_/H_e_), DXY = average absolute nucleotide divergence, F_ST_ = average genetic differentiation. †Calculated from AllSites dataset, *p<0.001. | Page 6 |
| **Table SB2.** Inferred mean migration rates with approximate 95% credible intervals estimated between subspecies (Antipodean and Gibson’s albatross) and islands (Antipodeans = Antipodes, Gibson’s = Adams, Disappointment) using BA3-SNPs with reads aligned to Gibson’s albatross reference genome. Approximate credible intervals were calculated as mean±1.96×SD per the BayeAss Edition 3.0 User’s Manual (Rannala, 2007). Proportion of non-migrants for each population shown along diagonal (light shading). | Page 6 |
| **Table SB3.** Cross-validation error from ADMIXTURE analyses with reads aligned to Gibson’s albatross reference genome. Cross-validation error was performed for K = 1-5. Lowest cross-validation error (shown in bold) represents the best predictive value of K (number of ancestral populations) for the dataset. | Page 7 |
| **Table SB4.** Contemporary effective population size (N_e_) of Antipodean and Gibson's albatross, calculated in NeEstimator2 with reads aligned to Gibson’s albatross reference genome. Upper and lower confidence intervals are based on jacknife resampling. | Page 7 |
| **Table SB5.** Genetic diversity statistics calculated using a dataset of 99 outlier SNPs for the Antipodean and Gibson’s albatross with reads aligned to Gibson’s albatross reference genome.  H_o_ = observed heterozysity (SE), H_e_ = expected heterozygosity (SE), F_IS_ = inbreeding coefficient (1 − H_o_/H_e_), F_ST_ = average genetic differentiation. *p<0.001 | Page 7 |
| **Table SB6.** Analysis of Molecular Variance (AMOVA) between Antipodean and Gibson's albatross populations using 524,726 neutral SNPs with reads aligned to Gibson’s albatross reference genome. | Page 8 |
| **Table SB7.** Analysis of Molecular Variance (AMOVA) between Antipodean and Gibson's albatross populations using 99 outlier SNPs with reads aligned to Gibson’s albatross reference genome. | Page 8 |

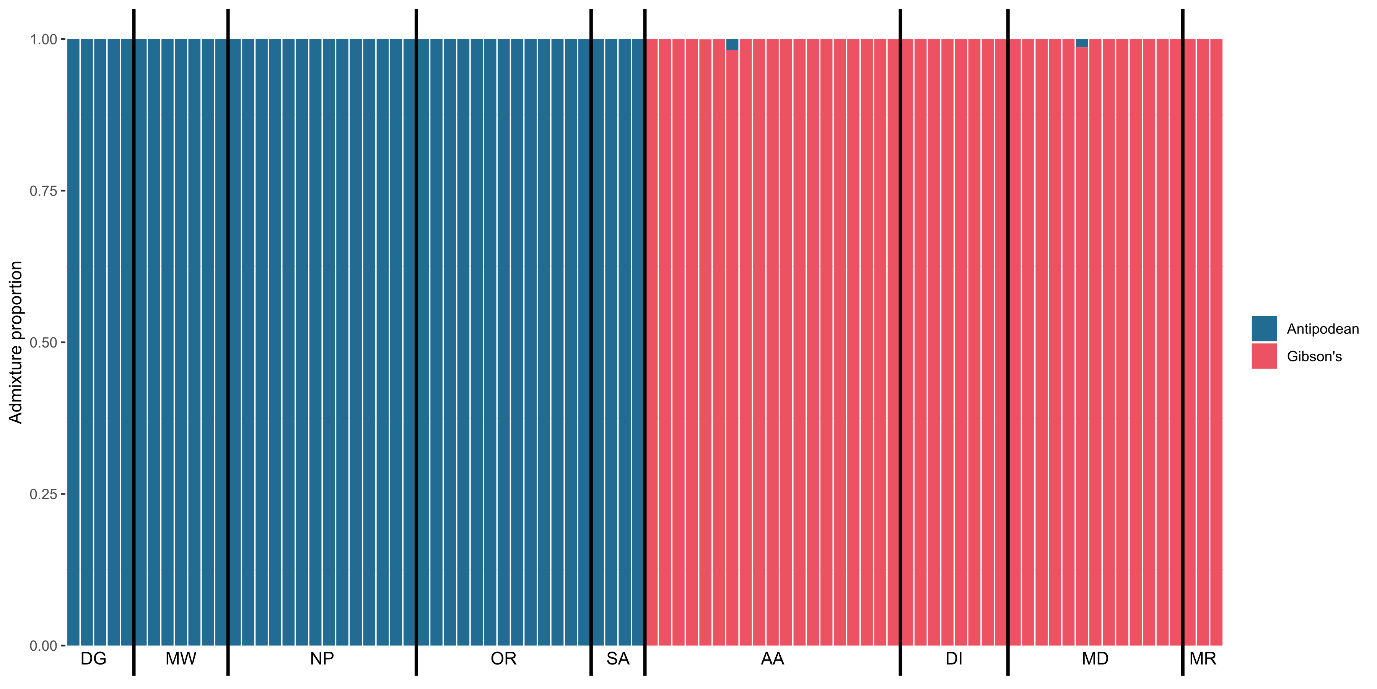

**Figure SB2**. ADMIXTURE plot for K = 2 showing the ancestry proportions of individuals from Antipodean and Gibson’s albatross populations using 524,726 neutral SNPs with reads aligned to Gibson’s albatross reference genome. Solid black lines indicate groupings of individuals from each sampling site. DG = Dougal, MW = Mt Waterhouse, NP = North Plains, OR = Orde Lees Stream, SA = Study Area, AA = Amherst Amsterdam, DI = Disappointment Island, MD = Mt Dick, MR = Maclaren Ridge

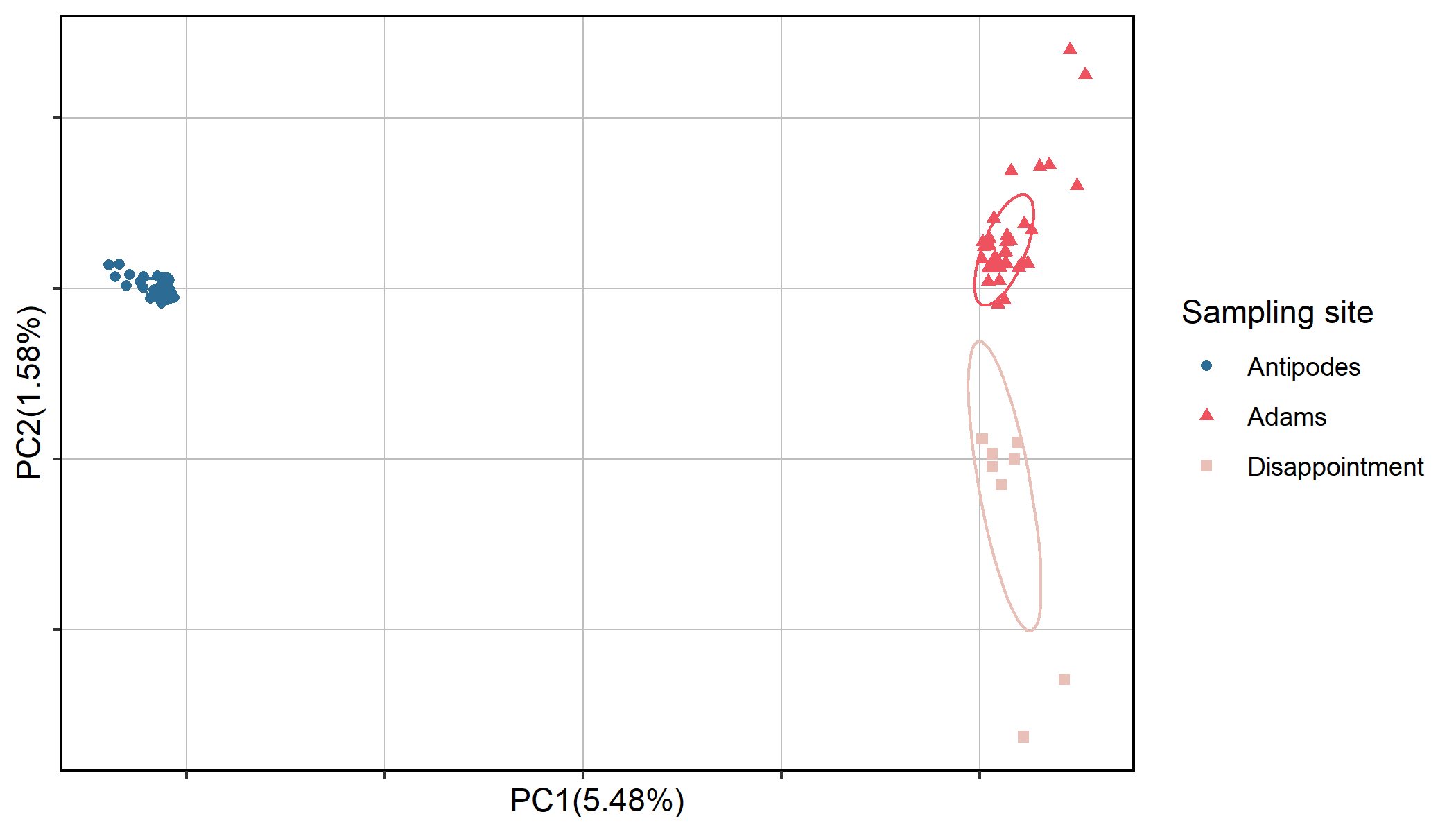

**Figure SB1.** Principal Component Analysis (PCA) based on 524,726 neutral SNPs showing the genetic clustering of Antipodean albatross (left cluster) and Gibson’s albatross (right cluster) with reads aligned to Gibson’s albatross reference genome. Samples are grouped by sampling island.

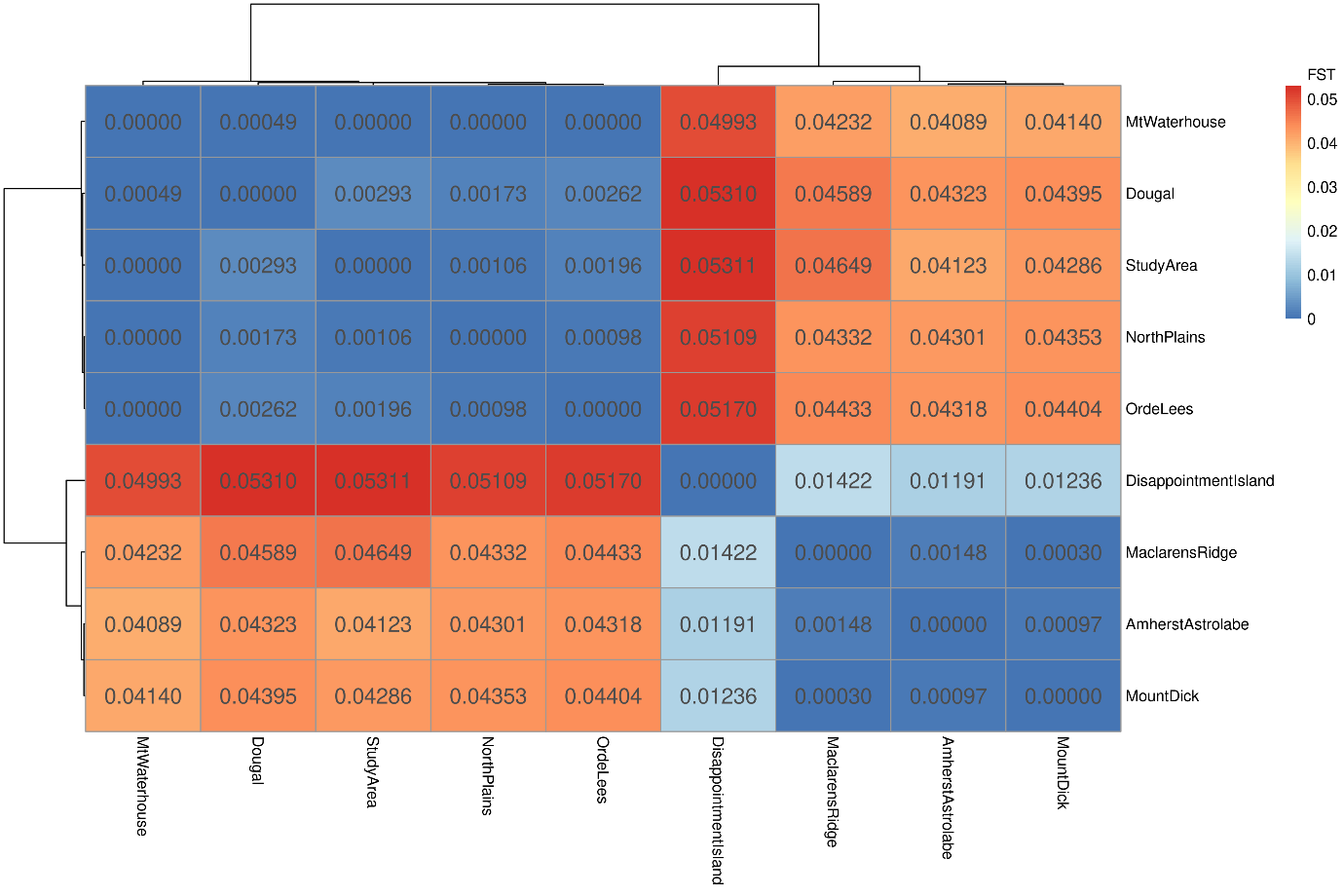

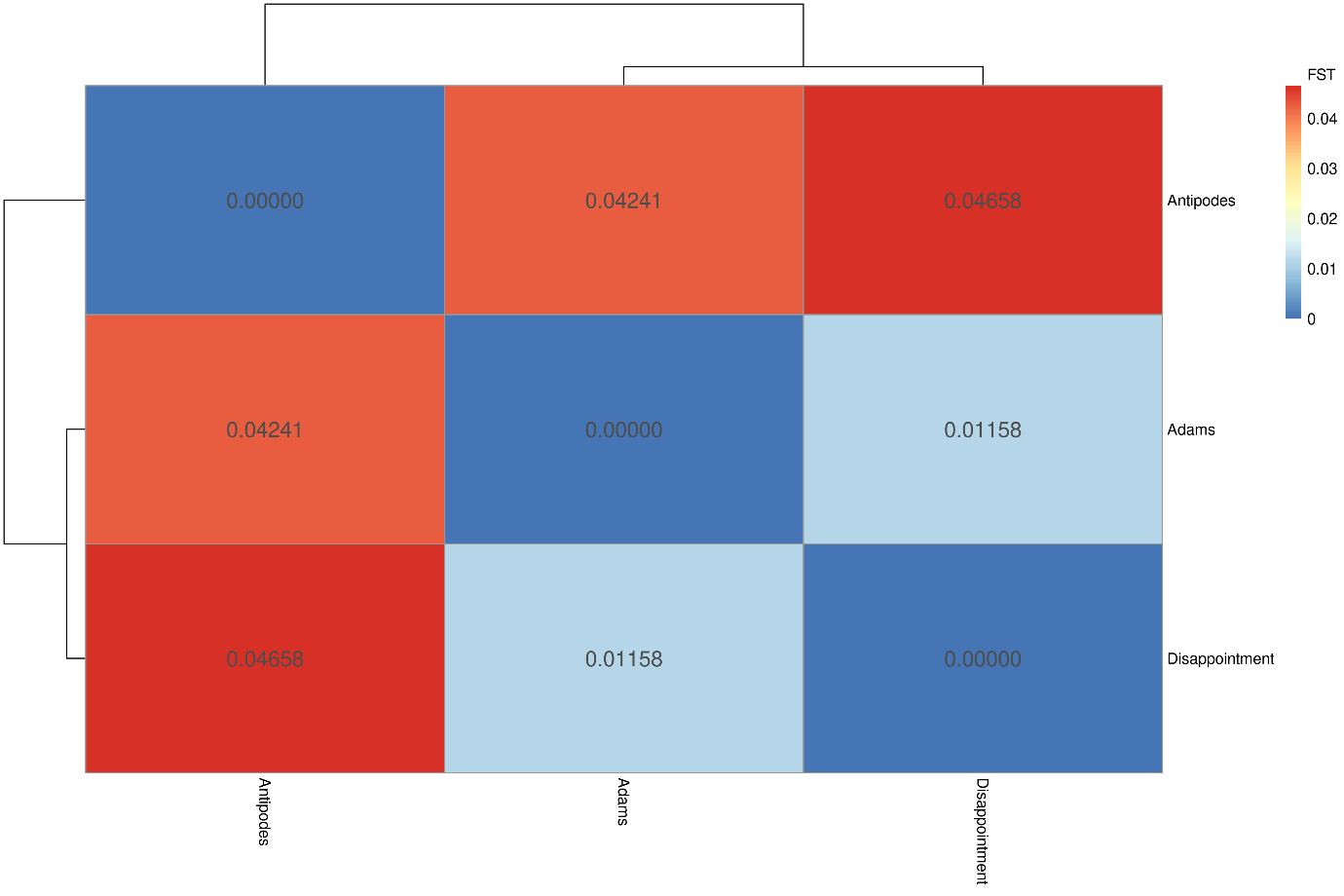

**A)**

**B)**

**Figure SB3.** Pairwise weighted F_ST_ estimates between **A)** all sampling sites and **B)** sampling islands of Antipodean and Gibson’s albatross with reads aligned to Gibson’s albatross reference genome.

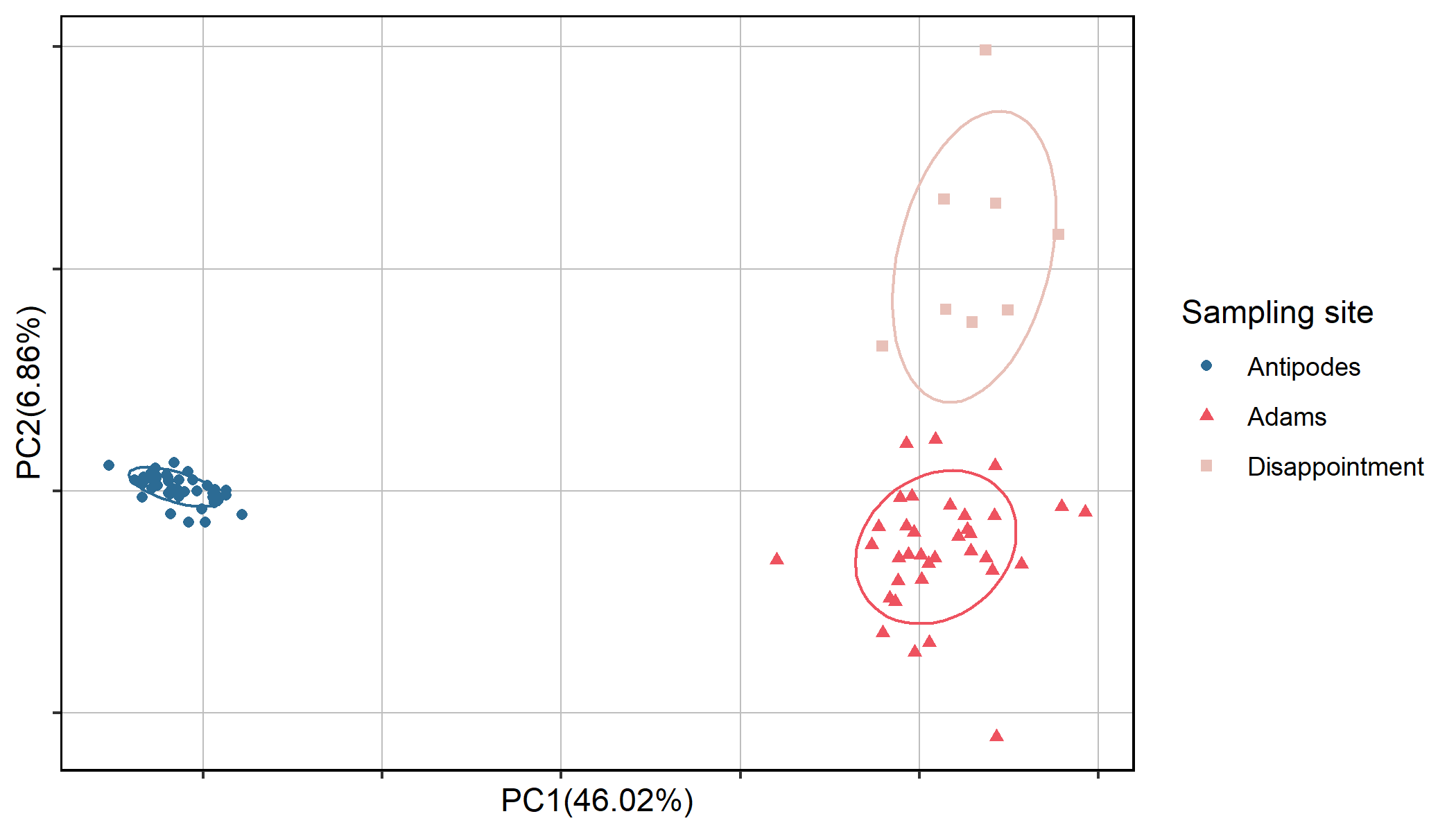

**Figure SB4.** Principal Components Analysis (PCA) based on 99 outlier SNPs showing the genetic clustering of Antipodean albatross (left cluster) and Gibson’s albatross (right cluster) with reads aligned to Gibson’s albatross reference genome. Samples are grouped by sampling island.

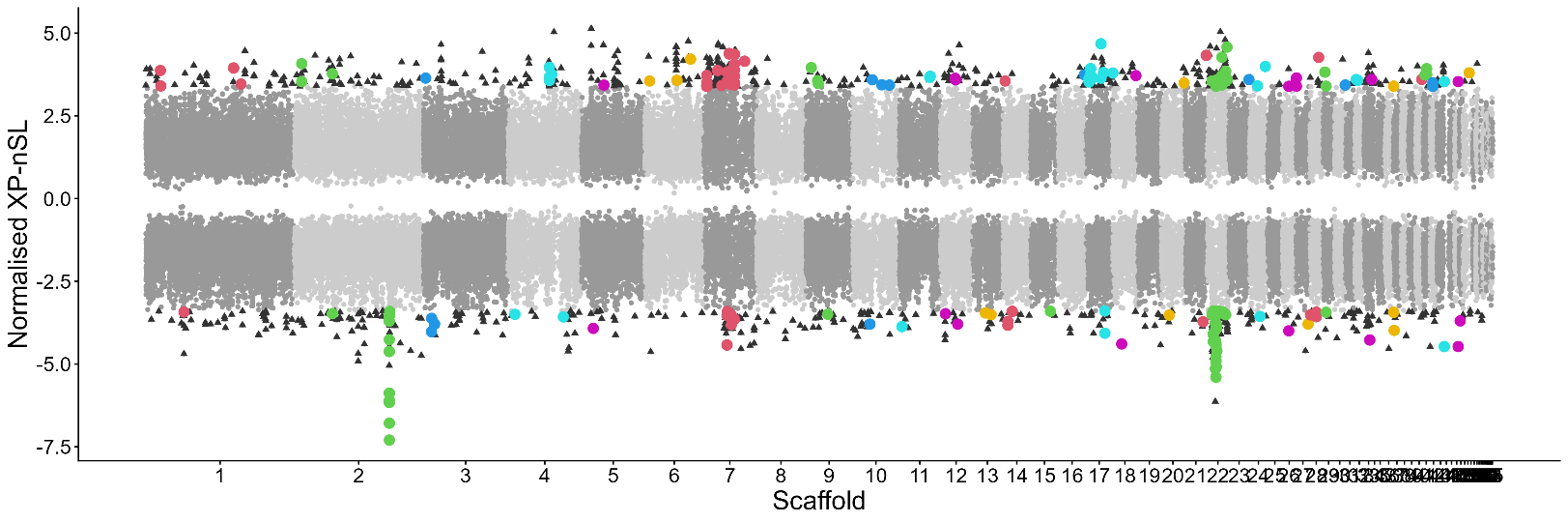

**Figure SB5.** Manhattan plot of normalised XP-nSL values in 10 kilobase (kb) windows across the genome in Antipodean and Gibson’s albatrosses with reads aligned to Gibson’s albatross reference genome. Positive and negative XP-nSL values suggest evidence for a selective sweep in the Antipodean and Gibson’s populations, respectively. Plot shows only the most extreme value per window (light and dark grey points, coloured by scaffold), with the top 1% most extreme values represented by black triangles. Windows within the top 1% that contain outlier SNPs identified by Baypass and pcadapt are shown by coloured circles (coloured by scaffold). Note: Scaffolds 2, 7 and 22 are homologous to scaffolds 7, 6 and 22 of the Antipodean albatross reference genome (Foote et al., 2026).

**Table SB1.** Genetic diversity statistics calculated using a dataset of 524,726 neutral SNPs or a genome-wide SNP dataset (including invariant sites) for the Antipodean and Gibson’s albatross with reads aligned to Gibson’s albatross reference genome.
*H*_o_ = observed heterozysity (SE), *H*_e_ = expected heterozygosity (SE), *π* = average nucleotide diversity, *θ* = average Watterson’s estimator, *F_IS_* = inbreeding coefficient (1 − *H*_o_/*H*_e_), *D_XY_* = average absolute nucleotide divergence, *F_ST_* = average genetic differentiation. †Calculated from AllSites dataset, *p<0.001.

|  | N | Ho | He | *π*† | *θ*† | *F_IS_* | *D_XY_*† | *F_ST_* |
| --- | --- | --- | --- | --- | --- | --- | --- | --- |
| Antipodean | 43 | 0.288 (0.0002) | 0.278 (0.0002) | 0.0008 | 0.0008 | -0.036 | 0.0009 | 0.04205* |
| Gibson’s | 43 | 0.292 (0.0002) | 0.280 (0.0002) | 0.0008 | 0.0008 | -0.043 |  |  |

**Table SB2.** Inferred mean migration rates with approximate 95% credible intervals estimated between subspecies (Antipodean and Gibson’s albatross) and islands (Antipodeans = Antipodes, Gibson’s = Adams, Disappointment) using BA3-SNPs with reads aligned to Gibson’s albatross reference genome. Approximate credible intervals were calculated as mean±1.96×SD per the BayeAss Edition 3.0 User’s Manual (Rannala, 2007). Proportion of non-migrants for each population shown along diagonal (light shading).

|  | *To* | Antipodean | Gibson’s | Antipodes | Adams | Dis. |
| --- | --- | --- | --- | --- | --- | --- |
| *From* | Antipodean | 0.9926 (0.9785 – 1.0067) | 0.0074 (-0.0067 – 0.0215) |  |  |  |
|  | Gibson’s | 0.0076 (-0.0069 – 0.0221) | 0.9924 (0.9779 – 1.0069) |  |  |  |
|  | Antipodes |  |  | 0.9855 (0.9660 – 1.0049) | 0.0087 (-0.0080 – 0.0254) | 0.0304  (-0.0241 – 0.0849) |
|  | Adams |  |  | 0.0072  (-0.0067 – 0.0211) | 0.9825 (0.9592 – 1.0058) | **0.2726 (0.1995 – 0.3457)** |
|  | Disappointment |  |  | 0.0073  (-0.0066 – 0.0212) | 0.0088  (-0.0079 – 0.0255) | 0.6970 (0.6427 – 0.7513) |

**Table SB3.** Cross-validation error from ADMIXTURE analyses with reads aligned to Gibson’s albatross reference genome. Cross-validation error was performed for K = 1-5. Lowest cross-validation error (shown in bold) represents the best predictive value of K (number of ancestral populations) for the dataset.

| K | CV error |
| --- | --- |
| 1 | 0.50701 |
| **2** | **0.49770** |
| 3 | 0.53158 |
| 4 | 0.56110 |
| 5 | 0.59701 |

**Table SB4.** Contemporary effective population size (N_e_) of Antipodean and Gibson's albatross, calculated in NeEstimator2 with reads aligned to Gibson’s albatross reference genome. Upper and lower confidence intervals are based on jacknife resampling.

| Population | N_e_ | CI for N_e_ | |
| --- | --- | --- | --- |
|  |  | Lower | Upper |
| Antipodean | 2,143 | 966 | ∞ |
| Gibson’s | 1,147 | 586 | 14,701 |

**Table SB5.** Genetic diversity statistics calculated using a dataset of 99 outlier SNPs for the Antipodean and Gibson’s albatross with reads aligned to Gibson’s albatross reference genome.
*H*_o_ = observed heterozysity (SE), *H*_e_ = expected heterozygosity (SE), *F_IS_* = inbreeding coefficient (1 − *H*_o_/*H*_e_), *F_ST_* = average genetic differentiation. *p<0.001

|  | N | Ho | He | *F_IS_* | *F_ST_* |
| --- | --- | --- | --- | --- | --- |
| Antipodean | 43 | 0.144 (0.0150) | 0.146 (0.0140) | 0.014 | 0.474* |
| Gibson’s | 43 | 0.403 (0.0140) | 0.394 (0.0094) | -0.023 |  |

**Table SB6.** Analysis of Molecular Variance (AMOVA) between Antipodean and Gibson's albatross populations using 524,726 neutral SNPs with reads aligned to Gibson’s albatross reference genome.

| Source of variation | d.f. | Sum of squares | Mean sum of squares | Variance components* | Percentage of variation | Φ | *p* |
| --- | --- | --- | --- | --- | --- | --- | --- |
| Between populations | 1 | 353,765.5 | 353,765.6 | 3,275.7 | 4.23 | 0.0162 | 0.001 |
| Between samples within populations | 84 | 6,052,496.7 | 72,053.5 | -2020.0 | -2.61 | -0.0273 | 0.889 |
| Within samples | 86 | 6,544,041.8 | 76093.5 | 76,093.5 | 98.38 | 0.0423 | 0.578 |
| **Total** | **171** | **12,950,304.2** | **75,732.7** | **77,349.2** | **100.00** |  |  |
| d.f. = degrees of freedom, *slightly negative covariance can occur in the absence of genetic structure, Φ = phi statistic, *p* = p-value. | | | | | | | |

**Table SB7.** Analysis of Molecular Variance (AMOVA) between Antipodean and Gibson's albatross populations using 99 outlier SNPs with reads aligned to Gibson’s albatross reference genome.

| Source of variation | d.f. | Sum of squares | Mean sum of squares | Variance components* | Percentage of variation | Φ | *p* |
| --- | --- | --- | --- | --- | --- | --- | --- |
| Between populations | 1 | 1,060.2 | 1,060.2 | 12.2 | 47.29 | 0.4729 | 0.001 |
| Between samples within populations | 84 | 1,136.6 | 13.5 | -0.03 | -0.13 | -0.0024 | 0.533 |
| Within samples | 86 | 1,169.3 | 13.6 | 13.6 | 52.83 | 0.4717 | 0.001 |
| **Total** | **171** | **3,366.1** | **19.7** | **25.7** | **100.00** |  |  |
| d.f. = degrees of freedom, *slightly negative covariance can occur in the absence of genetic structure, Φ = phi statistic, *p* = p-value. | | | | | | | |
